# Calcium dependence of neurotransmitter release at a high fidelity synapse

**DOI:** 10.1101/2021.05.15.444285

**Authors:** Abdelmoneim Eshra, Hartmut Schmidt, Jens Eilers, Stefan Hallermann

## Abstract

The Ca^2+^-dependence of the recruitment, priming, and fusion of synaptic vesicles are fundamental parameters controlling neurotransmitter release and synaptic plasticity. Despite intense efforts, these important steps in the synaptic vesicles’ cycle remain poorly understood because disentangling recruitment, priming, and fusion of vesicles is technically challenging. Here, we investigated the Ca^2+^-sensitivity of these steps at cerebellar mossy fiber synapses, which are characterized by fast vesicle recruitment mediating high-frequency signaling. We found that the basal free Ca^2+^ concentration (<200 nM) critically controls action potential-evoked release, indicating a high-affinity Ca^2+^ sensor for vesicle priming. Ca^2+^ uncaging experiments revealed a surprisingly shallow and non-saturating relationship between release rate and intracellular Ca^2+^ concentration up to 50 μM. Sustained vesicle recruitment was Ca^2+^-independent. Finally, quantitative mechanistic release schemes with five Ca^2+^ binding steps incorporating rapid vesicle recruitment via parallel or sequential vesicle pools could explain our data. We thus show that co-existing high and low-affinity Ca^2+^ sensors mediate recruitment, priming, and fusion of synaptic vesicles at a high-fidelity synapse.

## Introduction

During chemical synaptic transmission Ca^2+^ ions diffuse through voltage-gated Ca^2+^ channels, bind to Ca^2+^ sensors, and thereby trigger the fusion of neurotransmitter-filled vesicles (Südhof, 2012). The Ca^2+^-sensitivity of synaptic release is one of the most fundamental parameters influencing our understanding of fast neurotransmission. However, the Ca^2+^-sensitivity of the recruitment, priming, and fusion of synaptic vesicles is difficult to determine due to the large spatial gradients of the Ca^2+^ concentration, which occurs during Ca^2+^ influx through the Ca^2+^ channels. While the basal free intracellular Ca^2+^ concentration is ~50 nM, Ca^2+^ microdomains around the Ca^2+^ channels reach concentrations above 100 μM (Llinás et al., 1992). The technical development of caged Ca^2+^ compounds (Kaplan and Ellis-Davies, 1988) allows to experimentally elevate the Ca^2+^ concentration homogenously by photolysis and thus the direct measurement of the Ca^2+^-sensitivity of vesicle fusion (reviewed by Neher, 1998; Kochubey et al., 2011). First experiments with this technique at retinal bipolar cells of goldfish found a very low sensitivity of the release sensors with a half saturation at ~100 μM Ca^2+^ concentration and a fourth to fifth order relationship between Ca^2+^ concentration and neurotransmitter release (Heidelberger et al., 1994), similar to previous estimates at the squid giant synapse (Adler et al., 1991; Llinás et al., 1992). Subsequent work at other preparations showed different dose-response curves. For example, analysis of a central excitatory synapse, the calyx of Held (Forsythe, 1994) at a young pre-hearing age, found a much higher affinity with significant release below 5 μM intracellular Ca^2+^ concentration and similar slope of the dose-response curve (Bollmann et al., 2000; Lou et al., 2005; Schneggenburger and Neher, 2000; Sun et al., 2007). Further developmental analysis of the calyx of Held comparing the Ca^2+^-sensitivity of the release sensors at the age of P9 to P12-P15 (Kochubey et al., 2009) and P9 to P16-P19 (Wang et al., 2008) showed a developmental decrease in the Ca^2+^-sensitivity of vesicle fusion at the calyx of Held. A recent study at another excitatory central synapse, the hippocampal mossy fiber bouton, observed a high Ca^2+^-sensitivity of vesicle fusion in rather mature rats (P18–30; Fukaya et al., 2021), however the release rates in that study were not tested above 20 μM Ca^2+^ concentration. Analysis at an inhibitory central synapse revealed a high-affinity Ca^2+^ sensor and in addition a profoundly Ca^2+^-dependent priming step (Sakaba, 2008).

Moreover, analysis of the Ca^2+^-dependence of neurotransmitter release revealed a more shallow relationship between the rate of exocytosis and Ca^2+^ concentration at the sensory neurons of the rod photoreceptors (Duncan et al., 2010; Thoreson et al., 2004), and an absence of vesicle fusion below 7 μM Ca^2+^ concentration at the cochlear inner hair cells (Beutner et al., 2001).

Measuring the Ca^2+^-sensitivity of vesicle fusion is technically challenging and methodological errors could contribute to the differing Ca^2+^-sensitivity of various types of synapses. However, synapses show type-specific functional and structural differences (Atwood and Karunanithi, 2002; Nusser, 2018; Zhai and Bellen, 2004), which may lead to distinct Ca^2+^-sensitivities. Moreover, the rate at which new vesicles are recruited to empty release sites seems to be particularly different between synapses. The cerebellar mossy fiber bouton (cMFB) conveys high-frequency sensory information to the cerebellar cortex relying on extremely fast vesicle recruitment (Miki et al., 2020; Ritzau-Jost et al., 2014; Saviane and Silver, 2006). One aim of this study was therefore to determine the Ca^2+^-bsensitivity of vesicle fusion at mature cMFBs synapses at physiological temperature, and to test whether and how the prominent fast vesicle recruitment affects the Ca^2+^-mdependence of exocytosis at this synapse.

Compared with the Ca^2+^-sensitivity of vesicle fusion, the Ca^2+^-sensitivity of the vesicle recruitment and priming steps preceding fusion is even less well understood. While some studies at cMFBs proposed Ca^2+^-independent vesicle recruitment (Hallermann et al., 2010; Saviane and Silver, 2006), evidence for Ca^2+^-dependent steps preceding the fusion have been observed at several types of synapses (Awatramani et al., 2005; Doussau et al., 2017; Hosoi et al., 2007; Millar et al., 2005; Pan and Zucker, 2009; Sakaba, 2008). However, the dissection of vesicle recruitment, priming, and fusion is in general technically challenging. Therefore, we aimed to quantify the Ca^2+^-dependence of vesicle recruitment and priming at cMFBs by direct modification of the free intracellular Ca^2+^ concentration.

Our data revealed a strong dependence of the number of release-ready vesicles on basal Ca^2+^ concentrations between 30 and 180 nM, a significant release below 5 μM, and an apparent shallow dose-response curve in the studied Ca^2+^ concentration range of 1-50 μM. Computational simulations incorporating mechanistic release schemes with five Ca^2+^ binding steps and fast vesicle recruitment via sequential or parallel pools of vesicles could explain our data. Our results show the co-existence of Ca^2+^ sensors with high- and low-affinities that cover a large range of intracellular Ca^2+^ concentrations and mediate fast signaling at this synapse.

## Materials and Methods

### Preparation

Animals were treated in accordance with the German Protection of Animals Act and with the guidelines for the welfare of experimental animals issued by the European Communities Council Directive. Acute cerebellar slices were prepared from mature P35–P42 C57BL/6 mice of either sex as previously described (Hallermann et al., 2010). Isoflurane was used to anesthetize the mice which were then sacrificed by decapitation. The cerebellar vermis was quickly removed and mounted in a chamber filled with chilled extracellular solution. 300-μm-thick parasagittal slices were cut using a Leica VT1200 microtome (Leica Microsystems), transferred to an incubation chamber at 35 °C for ~30 min, and then stored at room temperature until use. The extracellular solution for slice cutting and storage contained (in mM) the following: NaCl 125, NaHCO_3_ 25, glucose 20, KCl 2.5, 2, NaH_2_PO_4_ 1.25, MgCl_2_ 1 (310 mOsm, pH 7.3 when bubbled with Carbogen [5% (vol/vol) O_2_/95% (vol/vol) CO_2_]). All recordings were restricted to lobules IV–V of the cerebellar vermis to reduce potential functional heterogeneity among different lobules (Straub et al., 2020).

### Presynaptic recordings and flash photolysis

All recordings were performed at physiological temperature by setting the temperature in the center of the recording chamber with immersed objective to 36°C using a TC-324B perfusion heat controller (Warner Instruments, Hamden, CT, United States). Presynaptic patch-pipettes were from pulled borosilicate glass (2.0/1.0 mm outer/inner diameter; Science Products) to open-tip resistances of 3-5 MΩ (when filled with intracellular solution) using a DMZ Puller (Zeitz-Instruments, Munich, Germany). Slices were superfused with artificial cerebrospinal fluid (ACSF) containing (in mM): NaCl 105, NaHCO_3_ 25, glucose 25, TEA 20, 4-AP 5, KCl 2.5, CaCl_2_ 2, NaH_2_PO_4_ 1.25, MgCl_2_ 1, and tetrodotoxin (TTX) 0.001, equilibrated with 95% O_2_ and 5% CO_2_. Cerebellar mossy fiber boutons (cMFBs) were visualized with oblique illumination and infrared optics (Ritzau-Jost et al., 2014). Whole-cell patch-clamp recordings of cMFBs were performed using a HEKA EPC10/2 amplifier controlled by Patchmaster software (HEKA Elektronik, Lambrecht, Germany). The intracellular solution contained (in mM): CsCl 130, MgCl_2_ 0.5, TEA-Cl 20, HEPES 20, Na2ATP 5, NaGTP 0.3. For Ca^2+^ uncaging experiments, equal concentrations of DM-nitrophen (DMn) and CaCl_2_ were added depending on the aimed post-flash Ca^2+^ concentration, such that either 0.5, 2, or 10 mM was used for low, middle, or high target range of post-flash Ca^2+^ concentration, respectively (Supplementary Table 1). To quantify post-flash Ca^2+^ concentration with a previously established dual indicator method (see below; Delvendahl et al., 2015; Sabatini et al., 2002), Atto594, OGB-5N, and Fluo-5F were used at concentrations as shown in (Supplementary Table 1).

**Supplementary Table 1.** Parameters for weak, middle, and strong post-flash Ca^2+^ elevations.

|  | **weak Ca<sup>2+</sup>**<br>**elevation** | **middle Ca<sup>2+</sup>**<br>**elevation** | **strong Ca<sup>2+</sup>**<br>**elevation** |
| --- | --- | --- | --- |
| **UV illumination** |  |  |  |
| Duration (ms) | 0.1 or 1 | 0.1 | 0.1 or 0.2 |
| Intensity (%) | 10 - 100 | 20 - 100 | 100 |
| **Concentration in intracellular solution (mM)** |  |  |  |
| ATTO 594 | 0.010 | 0.020 | 0.020 |
| Fluo 5F | 0.050 | 0 | 0 |
| OGB 5N | 0 | 0.200 | 0.200 |
| CaCl<sub>2</sub> | 0.500 | 2.000 | 10.000 |
| DM-N | 0.500 | 2.000 | 10.000 |
| **Obtained peak post-flash Ca<sup>2+</sup> (μM)** |  |  |  |
| Min | 1.1 | 2.7 | 15.7 |
| Max | 7.1 | 36.0 | 62.6 |
| Median | 2.4 | 8.8 | 25.1 |
| **Simulated uncaging fraction of DMn** |  |  |  |
| $\alpha$ | 0.08-0.5 | 0.15-0.55 | 0.14-0.25 |

A 50 mM solution stock of DMn was prepared by neutralizing 50 mM DMn in H_2_O with 200 mM CsOH in H_2_O. The purity of each DMn batch was determined in the intracellular solution used for patching through titration with sequential addition of Ca^2+^ as previously described (Schneggenburger, 2005) and by measuring the Ca^2+^ concentration using the dual indicator method with 10 μM Atto594 and 50 μM OGB1 (Delvendahl et al., 2015).

After waiting for at least one minute in whole-cell mode to homogenously load the terminal with intracellular solution, capacitance measurements were performed at a holding potential of −100 mV with sine-wave stimulation (5 kHz or 10 kHz frequency and ±50 mV amplitude; Hallermann et al., 2003). During the ongoing sine-wave stimulation, a UV laser source (375 nm, 200 mW, Rapp OptoElectronic) was used to illuminate the whole presynaptic terminal. According to a critical illumination, the end of the light guide of the UV laser was imaged into the focal plan resulting in a homogeneous illumination in a circular area of ~30 μm diameter (Fig. 2 – figure supplement 1). The duration of the UV illumination was 100 μs controlled with sub-microsecond precision by an external triggering of the laser source. In capacitance measurements with 10 kHz sine wave frequency, longer pulses of 200 μs were used to reach high Ca^2+^ levels. In a subset of experiments, UV pulses of 1 ms were used to rule out fast undetectable Ca^2+^ overshoots (Bollmann et al., 2000; Fig. 3 – figure supplement 3). The UV flash intensity was set to 100% and reduced in some experiments (10 – 100%) to obtain small elevations in Ca^2+^ concentrations (Supplementary Table 1). All chemicals were from Sigma-Aldrich. Atto594 was purchased from Atto-Tec, Ca^2+^-sensitive fluorophores from Life Technologies, and DMn from Synaptic Systems.

### Paired Recordings between cMFBs and GCs

For paired pre- and postsynaptic recordings, granule cells (GCs) were whole-cell voltage-clamped with intracellular solution containing the following (in mM): K-gluconate 150, NaCl 10, K-HEPES 10, MgATP 3 and Na-GTP 0.3 (300–305 mOsm, pH adjusted to 7.3 with KOH). 10 μM Atto594 was included to visualize the dendrites of the GCs (Ritzau-Jost et al., 2014). After waiting sufficient time to allow for the loading of the dye, the GC dendritic claws were visualized through two-photon microscopy, and subsequently, cMFBs near the dendrites were identified by infrared oblique illumination and were patched and loaded with caged Ca^2+^ and fluorescent indicators as previously described. The reliable induction of an EPSC in the GC was used to unequivocally confirm a cMFB-GC synaptic connection. In a subset of the Ca^2+^ uncaging experiments, simultaneous presynaptic capacitance and postsynaptic EPSC recordings were performed from GC and cMFB, respectively.

### Clamping intracellular basal Ca^2+^ concentrations

The intracellular solution for presynaptic recordings of the data shown in Fig. 1 contained the following in mM: K-gluconate 150, NaCl 10, K-HEPES 10, MgATP 3, Na-GTP 0.3. With a combination of EGTA and CaCl_2_ (5 mM EGTA / 0.412 mM CaCl_2_ or 6.24 mM EGTA / 1.65 mM CaCl_2_), we aimed to clamp the free Ca^2+^ concentration to low and high resting Ca^2+^ concentrations of ~50 or ~200 nM, respectively, while maintaining a free EGTA concentration constant at 4.47 mM. The underlying calculations were based on a Ca^2+^ affinity of EGTA of 543 nM (Lin et al., 2017). The resulting free Ca^2+^ concentration was quantified with the dual indicator method (see below) and was found to be to ~30 or ~180 nM, respectively (Fig. 1A).

**Figure 1.**
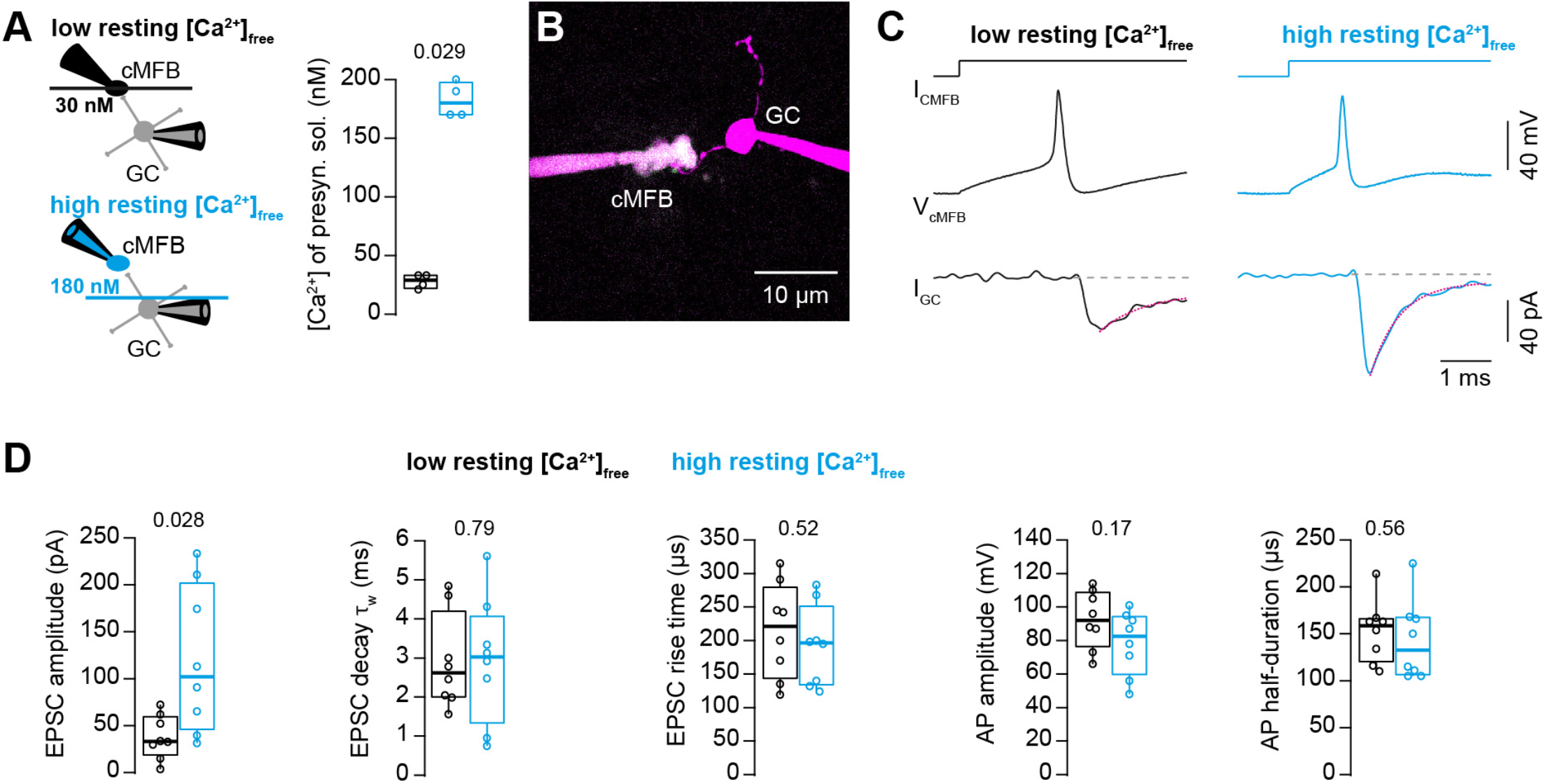
Action potential-evoked synaptic release critically depends on basal intracellular Ca^2+^ concentration. A. *Left:* Illustration of the cellular connectivity of the cMFB to GC synapse during simultaneous pre- and postsynaptic patch-clamp recording. The presynaptic terminal was loaded with an intracellular solution having either low or high free basal Ca^2+^ concentration (top and bottom, respectively). *Right:* Comparison of the average free Ca^2+^ concentration in the presynaptic patch pipette (quantified by two-photon Ca^2+^ imaging) for the intracellular solutions with low and high basal Ca^2+^ (n = 4 each). B. Example two-photon microscopic image of a cMFB and a GC in the paired whole-cell configuration. C. Example traces of a paired cMFB-GC recording with current injection (I_cMFB_) (*top*) eliciting an action potential in the cMFB (*middle*) and an EPSC in the postsynaptic GC (*bottom*). Black and blue color code corresponds to low and high free basal Ca^2+^ concentration in the presynaptic solution, respectively. The decay of the EPSC was fitted with a bi-exponential function (magenta line). D. Comparison of the properties of presynaptic action potentials and EPSCs evoked after eliciting an action potential in the presynaptic terminal using solutions having either low (black) or high (blue) free Ca^2+^ concentration. From left to right: peak amplitude of the EPSC, weighted decay time constant of the EPSC, 10-to-90% rise time of the EPSC, amplitude of the presynaptic action potential, and action potential half-duration (n = 8 and 8 pairs for the conditions with low and high resting Ca^2+^ concentration, respectively).

### Quantitative two-photon Ca^2+^ imaging

For the quantification of Ca^2+^ signals elicited through UV flash-induced uncaging, two-photon Ca^2+^ imaging was performed as previously described (Delvendahl et al., 2015) using a Femto2D laser-scanning microscope (Femtonics) equipped with a pulsed Ti:Sapphire laser (MaiTai, SpectraPhysics) adjusted to 810 nm, a 60×/1.0 NA objective (Olympus), and a 1.4 NA oil-immersion condenser (Olympus). Data were acquired by doing line-scans through the cMFB. To correct for the flash-evoked luminescence from the optics, the average of the fluorescence from the line-scan in an area outside of the bouton was subtracted from the average of the fluorescence within the bouton (Fig. 2B). Imaging data were acquired and processed using MES software (Femtonics). Upon releasing Ca^2+^ from the cage, we measured the increase in the green fluorescence signal of the Ca^2+^ sensitive indicator (OGB-5N or Fluo-5F) and divided it by the fluorescence of the Ca^2+^ insensitive Atto594 (red signal). The ratio (R) of green-over-red fluorescence was translated into a Ca^2+^ concentration through the following calculation (Yasuda et al., 2004).

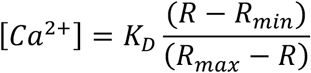

**Figure 2.**
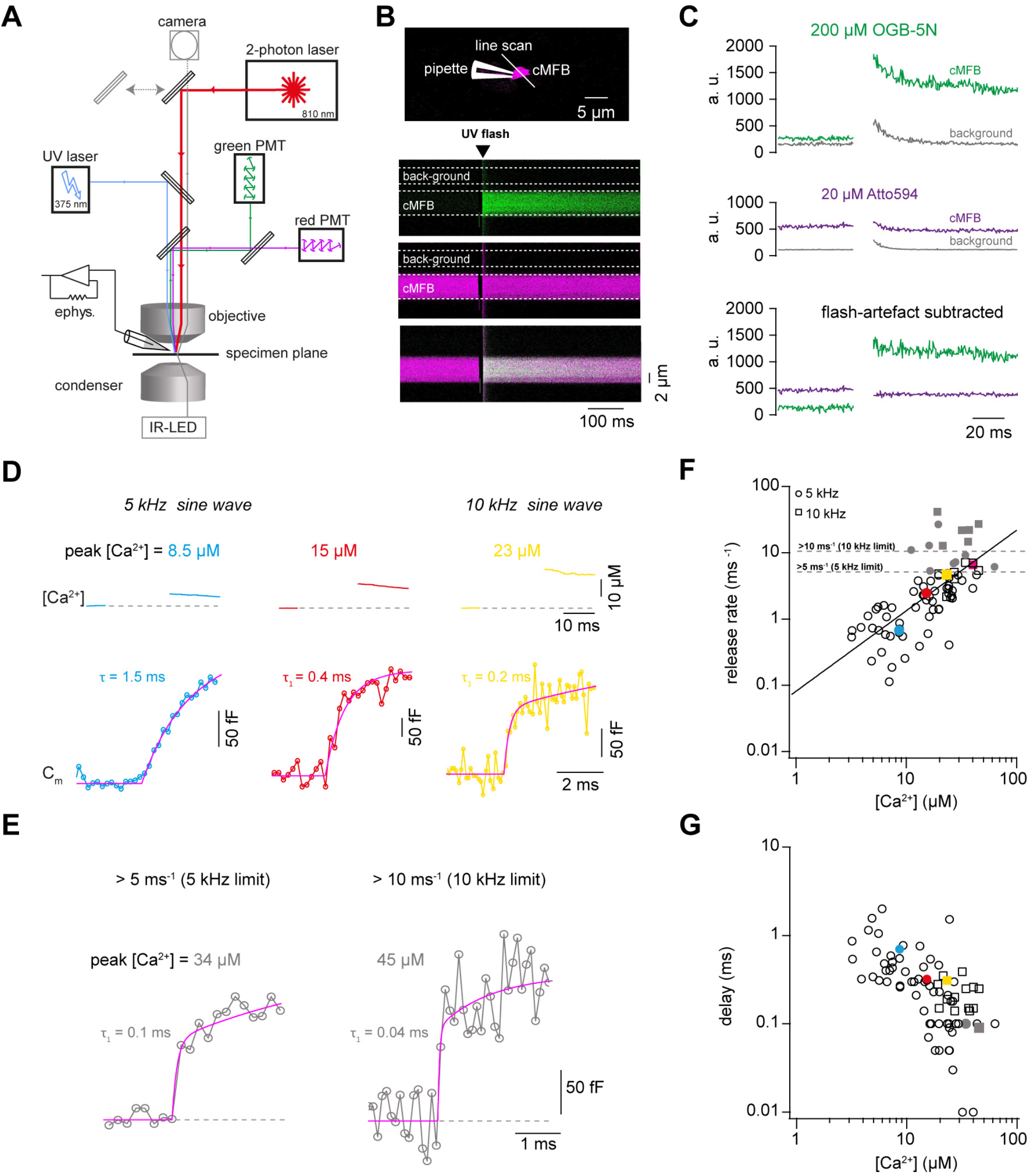
Ca^2+^ uncaging dose-response curve measured with presynaptic capacitance measurements. A. Illustration of the experimental setup showing the light path of the two-photon laser illumination (red line), the UV laser illumination (blue line), the electrophysiology amplifier (‘ephys.’), the red and green gate-able photomultiplier tubes (PMTs), and infrared LED illumination with oblique illumination via the condenser for visualization of the cells at the specimen plane by the camera (grey line) when the upper mirror is moved out of the light path (grey arrow). B. *Top:* Two-photon microscopic image of a cMFB in the whole-cell configuration loaded with OGB-5N, Atto594, and DMn/ Ca^2+^. Positions of the patch pipette and line scan are indicated. *Bottom:* Two-photon line scan showing the fluorescence signal as measured through the green PMT, red PMT, and an overlay of the green and red channels. Arrow indicates the onset of the UV flash and dashed lines represent the flash-induced luminescence artefact as detected outside the cMFB. The lookup tables for the green and red channel were arbitrarily but linearly adjusted independent of the absolute values in C. C. *Top:* change in fluorescence intensity within the cMFB for the green channel along with the corresponding flash-induced green artefact measured in the background. *Middle:* change in fluorescence intensity within the cMFB for the red channel along with the corresponding flash-induced red artefact. *Bottom:* green and red fluorescence signal after subtracting the flash-induced artefacts. D. *Top:* Ca^2+^ signals of different concentrations elicited through Ca^2+^ uncaging in three different cells, the flash was blanked. *Bottom:* corresponding traces of capacitance recordings measured using a 5 kHz sinusoidal stimulation (left and middle) or 10 kHz sinusoidal stimulation (right). *τ* represents the time constant from a mono-exponential fit, *τ_1_* represents the time constant of the fast component of a bi-exponential fit. E. Traces of capacitance recordings showing the resolution limit in detecting fast release rates of >5 ms^-1^ using 5 kHz sinusoidal stimulation or >10 ms^-1^ using 10 kHz sinusoidal stimulation. F. Plot of release rate versus post-flash Ca^2+^ concentration. The line represents a fit with a Hill equation (eq. 2) with best-fit values *V_max_* = 1.7*10^7^ ms^-1^, *K_D_* = 7.2*10^6^ μM, and *n* = 1.2. Color coded symbols correspond to traces in D – E. Grey symbols represent values above the resolution limit. G. Plot of synaptic delay versus post-flash Ca^2+^ concentration. Color coded symbols correspond to traces in D – E.

To avoid pipetting irregularities, which might influence the quantification of the fluorescence signals, pre-stocks of Ca^2+^-sensitive and Ca^2+^-insensitive indicators were used. For each pre-stock and each intracellular solution, 10 mM EGTA or 10 mM CaCl_2_ were added to measure minimum (R_min_) and maximum (R_max_) fluorescence ratios, respectively. We performed these measurements in cMFBs and GCs as well as in cuvettes. Consistent with a previous report (Delvendahl et al., 2015), both R_min_ and R_max_ were higher when measured in cells than in cuvettes (by a factor of 1.73 ± 0.05; n = 83 and 63 measurements in situ and in cuvette; Fig. 3 – figure supplement 2A). The values in cMFBs and GCs were similar (Fig. 3 – figure supplement 2B). OGB-5N is not sensitive in detecting Ca^2+^ concentrations less than 1 μM. Therefore, we deliberately adjusted R_min_ of OGB-5N in the recordings where the pre-flash Ca^2+^ had negative values, to a value resulting in a pre-flash Ca^2+^ concentration of 60 nM, which corresponds to the average resting Ca^2+^ concentration in these boutons (Delvendahl et al., 2015). This adjustment of R_min_ resulted in a reduction of post-flash Ca^2+^ amplitudes of on average 7.5 ± 0.4 % (n = 37). Without this adjustment, the estimated *K_D_* of the Ca^2+^ sensors for release would be even slightly higher.

The fluorescence properties of DMn change after flash photolysis, and the Ca^2+^ sensitive and insensitive dyes can differentially bleach during UV flash (Schneggenburger, 2005; Zucker, 1992). We assumed no effect of the UV flash on the *K_D_* of the Ca^2+^ sensitive dyes (Escobar et al., 1997), and measured R_min_ and R_max_ before and after the flash for each used UV flash intensity and duration in each of the three solutions (Supplementary Table 1; Schneggenburger et al., 2000). The flash-induced change was strongest for R_max_ of solutions with OGB-5N, but reached only ~20% with the strongest flashes (Fig. 3 – figure supplement 2C – 2F).

### Deconvolution

Deconvolution of postsynaptic currents was performed essentially as described by Ritzau-Jost et al. (2014), based on routines developed by Neher and Sakaba (2001b). The principle of this method is that the EPSC comprises currents induced by synchronous release and residual glutamate in the synaptic cleft due to delayed glutamate clearance and glutamate spill-over from neighboring synapses. Kynurenic acid (2 mM) and Cyclothiazide (100 μM) were added to the extracellular solution to reduce postsynaptic receptor saturation and desensitization, respectively. The amplitude of the miniature EPSC (mEPSC) was set to the mean value of 10.1 pA (10.1 ± 0.2 pA; n = 8) as measured in 2 mM kynurenic acid and 100 μM cyclothiazide.

The deconvolution kernel had the following free parameters: the mEPSC early slope τ_0_, the fractional amplitude of the slow mEPSC decay phase α, the time constant of the slow component of the decay τ_2_ of the mEPSC, the residual current weighting factor β, and the diffusional coefficient *d*. Applying the “fitting protocol” described by Neher and Sakaba (2001b) before flash experiments might affect the number of vesicles released by subsequent Ca^2+^ uncaging. On the other hand, applying the “fitting protocol” after Ca^2+^ uncaging might overestimate the measured number of vesicles due to flash-induced toxicity and synaptic fatigue especially when applying strong Ca^2+^ uncaging. Therefore, we used the experiments with weak and strong flashes to extract the mini-parameters and the parameters for the residual current, respectively, as described in the following in more detail. To obtain the mini parameters (early slope, α, and τ_2_) using weak flashes, deconvolution was first performed with a set of trial parameters for each cell pair. The mini-parameters of the deconvolution were optimized in each individual recording to yield low (but non-negative) step-like elevations in the cumulative release corresponding to small EPSCs measured from the postsynaptic terminal (the parameters for the residual current had little impact on the early phase of the cumulative release rate within the first 5 ms, therefore, some reasonable default values for the parameters of the residual current were used while iteratively adjusting the fast mini parameters for each individual recording). Next, using the average of the mini-parameters obtained from weak flashes, the deconvolution parameters for the residual current (β and d) were optimized in each recording with strong flashes until no drops occurred in the cumulative release in the range of 5 – 50 ms after the stimulus (while iteratively readjusting the mini parameters, if needed, to avoid any drops in the cumulative release in the window of 5 – 10 ms that might arise when adjusting the slow parameters based on the cumulative release in the range of 5 – 50 ms). Finally, we averaged the values of each parameter and the deconvolution analysis of all recordings was re-done using the average parameters values. To test the validity of this approach, cumulative release from deconvolution of EPSCs and presynaptic capacitance recordings were compared in a subset of paired recordings (Ritzau-Jost et al., 2014). Exponential fits to the cumulative release and the presynaptic capacitance traces provided average time constants of 2.43 ± 0.81 and 2.65 ± 0.88 ms, respectively (n = 9 pairs). On a paired-wise comparison, the difference in the time constant was always less than 40%. Therefore, both approaches yielded similar results.

To measure the number of GCs connected by one cMFB, we compared the product of the amplitude and the inverse of the time constant of the exponential fits of presynaptic capacitance trace and the simultaneously measured cumulative release trace obtained by deconvolution analysis of EPSC. Assuming a capacitance of 70 aF per vesicle (Hallermann 2003), we obtained an average value of 90.1 GCs per MFB in close agreement with previous estimates using a similar approach (Ritzau-Jost et al., 2014). This connectivity ratio is larger than previous estimates (~10, Billings et al., 2014; ~50, Jakab and Hamori, 1988) which could be due to a bias towards larger terminals, ectopic vesicle release, postsynaptic rundown, or release onto Golgi cells.

### Measurement of Ca^2+^ concentration using a Ca^2+^-sensitive electrode

A precise estimation of the binding affinity of the Ca^2+^ sensitive dyes is critical in translating the fluorescence signals into Ca^2^ concentration. It has been reported that the *K_D_* of fluorescent indicators differs significantly depending on the solution in which it is measured (Tran et al., 2018) due to potential differences in ionic strength, pH, and concentration of other cations. Accordingly, different studies have reported different estimates of the *K_D_* of OGB-5N having an up to 8-fold variability (Delvendahl et al., 2015; Digregorio and Vergara, 1997; Neef et al., 2018). In these studies, the estimation of the *K_D_* of the Ca^2+^ sensitive dyes depended on the estimated *K_D_* of the used Ca^2+^ chelator, which differs based on the ionic strength, pH, and temperature of the solution used for calibration. So, we set out to measure the K_D_ of OGB-5N, in the exact solution and temperature which we used during patching, through direct potentiometry using an ion-selective electrode combined with two-photon Ca^2+^ imaging. An ion-selective electrode for Ca^2+^ ions provides a direct readout of the free Ca^2+^ concentration independent of the *K_D_* of the used Ca^2+^ chelator. Using the same intracellular solution and temperature as used during experiments, the potential difference between the Ca^2+^-sensitive electrode (ELIT 8041 PVC membrane, NICO 2000) and a single junction silver chloride reference electrode (ELIT 001n, NIC0 2000) was read out with a pH meter in mV mode. A series of standard solutions, with defined Ca^2+^ concentration (Thermo Fisher) covering the whole range of our samples, were used to plot a calibration curve of the potential (mV) versus Ca^2+^ concentration (μM). Then, the potential of several sample solutions containing the same intracellular solution used for patching, but with different Ca^2^ concentrations buffered with EGTA, was determined. This way, we got a direct measure of the free Ca^2+^ concentration of several sample solutions, which were later used after the addition of Ca^2+^ sensitive fluorometric indicators to plot the fluorescence signal of each solution versus the corresponding free Ca^2+^ concentration verified by the Ca^2+^-bsensitive electrode, and accordingly the *K_D_* of the Ca^2+^ indicators were obtained from fits with Hill equation. The estimated *K_D_* was two-fold higher than the estimate obtained using only the Ca^2+^ Calibration Buffer Kit (Thermofischer) without including intracellular patching solution (Fig. 3 – figure supplement 1). Comparable results were obtained when estimating the free Ca^2+^ concentration using Maxchelator software (https://somapp.ucdmc.ucdavis.edu/pharmacology/bers/maxchelator/). Therefore, we used two independent approaches to confirm the *K_D_* of OGB-5N. We found that TEA increased the potential of the solutions measured through the Ca^2+^-sensitive electrode, which is consistent with a previous report showing a similar effect of quaternary ammonium ions on potassium sensitive microelectrodes (Neher and Lux, 1973). We compared the fluorescence signals of our samples with or without TEA, to check if this effect of TEA is due to an interaction with the electrode or due to an effect on the free Ca^2+^ concentration, and found no difference. Therefore, TEA had an effect on the electrode read-out without affecting the free Ca^2+^, and accordingly, TEA was removed during the potentiometric measurements (Fig. 3 – figure supplement 1). This resulted in a good agreement of the estimates of the free Ca^2+^ concentration measured using a Ca^2+^-sensitive electrode and those calculated via Maxchelator.

### Assessment of the UV energy profile

The homogeneity of the UV laser beam at the specimen plane was assessed *in vitro* by uncaging fluorescein (CMNB-caged fluorescein, Invitrogen). Caged fluorescein (2 mM) was mixed with glycerol (5% caged fluorescein/ 95% glycerol) to limit the mobility of the released dye (Bollmann et al., 2000). We did the measurements at the same plane as we put the slice during an experiment. The fluorescence profile of the dye after being released from the cage was measured at different z-positions over a range of 20 μm. The intensity of fluorescein was homogenous over an area of 10 μm x 10 μm which encompasses the cMFB.

### Data analysis

The increase in membrane capacitance and in cumulative release based on deconvolution analysis was fitted with the following single or bi-exponential functions using Igor Pro (WaveMetrics) including a baseline and a variable onset.

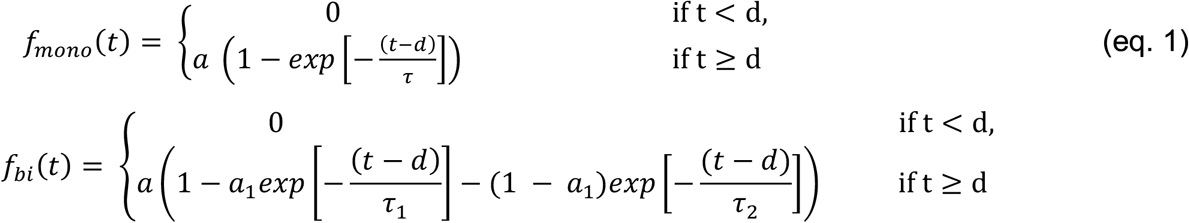

where *d* defines the delay, *a* the amplitude, *τ* the time constant of the mono-exponetial fit, τ_1_, and τ_2_ the time constant of the fast and slow component of the bi-exponential fit, respectively, and *a*_1_ the relative contribution of the fast component of the bi-exponential fit. The fitting of the release traces was always done with a time window of 5 ms before and 10 ms after flash onset. If the time constant of the mono-exponential fit exceeded 10 ms, a longer fitting duration of 60 ms after flash onset was used.

The acceptance of a bi-exponential fit was based on the fulfillment of the following three criteria: (1) at least 4% decrease in the sum of squared differences between the experimental trace and the fit compared with a mono-exponential fit (χ^2^mono/χ^2^bi > 1.04), (2) the time constants of the fast and the slow components differed by a factor >3, and (3) the relative contribution of each component was >10% (i.e. 0.1 < *a*_1_ < 0.9). If any of these criteria was not met, a mono-exponential function was used instead. In the case of weak flashes, where we could observe single quantal events within the initial part of the EPSC, mono-exponential fits were applied. In Fig. 1, bi-exponential functions were used to fit the decay of the EPSC and the weighted time constants were used.

Hill equations were used to fit the release rate versus intracellular Ca^2+^ concentration on a double logarithmic plot according to the following equation:

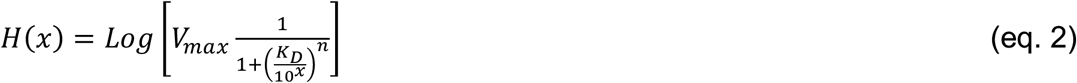

where *Log* is the decadic logarithm, *V_max_* the maximal release rate, *K_D_* the Ca^2+^ concentration at the half-maximal release rate, and *n* the Hill coefficient. H(x) was fit on the decadic logarithm of the release rates and x was the decadic logarithm of the intracellular Ca^2+^ concentration.

### Modeling of intra-bouton Ca^2+^ dynamics

We simulated the intra-bouton Ca^2+^ dynamics using a single compartment model. The kinetic reaction schemes for Ca^2+^ and Mg^2+^ uncaging and -binding (Fig. 6A) were converted to a system of ordinary differential equations (ODEs) that was numerically solved using the NDSolve function in Mathematica 12 (Wolfram) as described previously (Bornschein et al., 2019). The initial conditions for the uncaging simulation were derived by first solving the system of ODEs for the steady state using total concentrations of all species and the experimentally determined [Ca^2+^]_rest_ as starting values. Subsequently, the values obtained for all free and bound species were used as initial conditions for the uncaging simulation. The kinetic properties of DMn were simulated according to Faas et al. (2005, 2007). The total DMn concentration ([DMn]_T_) includes the free form ([DMn]), the Ca^2+^ bound form ([CaDMn]), and the Mg^2+^ bound form ([MgDMn]). Each of these forms is subdivided into an uncaging fraction (α) and a non-uncaging fraction (1-α). The uncaging fraction were further subdivided into a fast (af) and a slow (1-af) uncaging fraction:

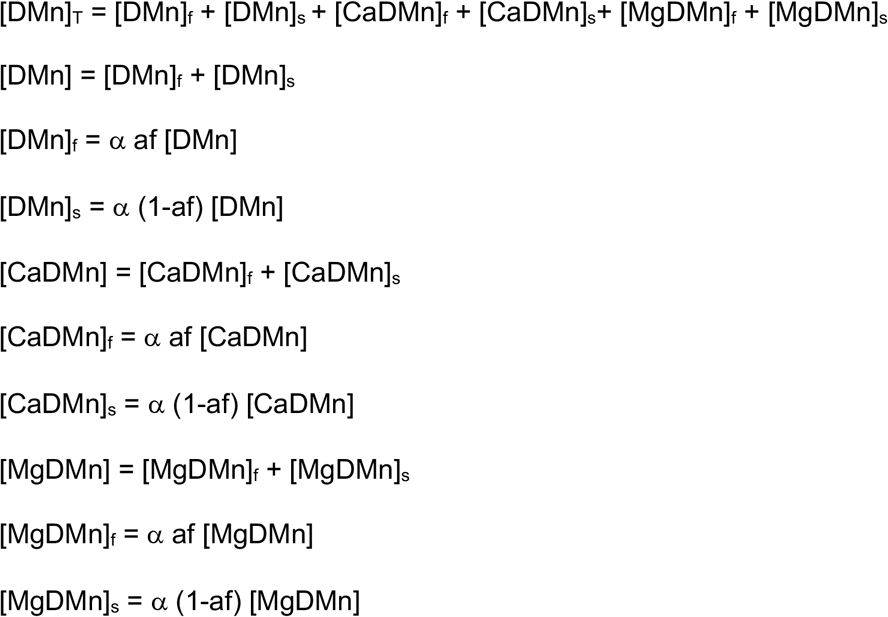

**Figure 3.**
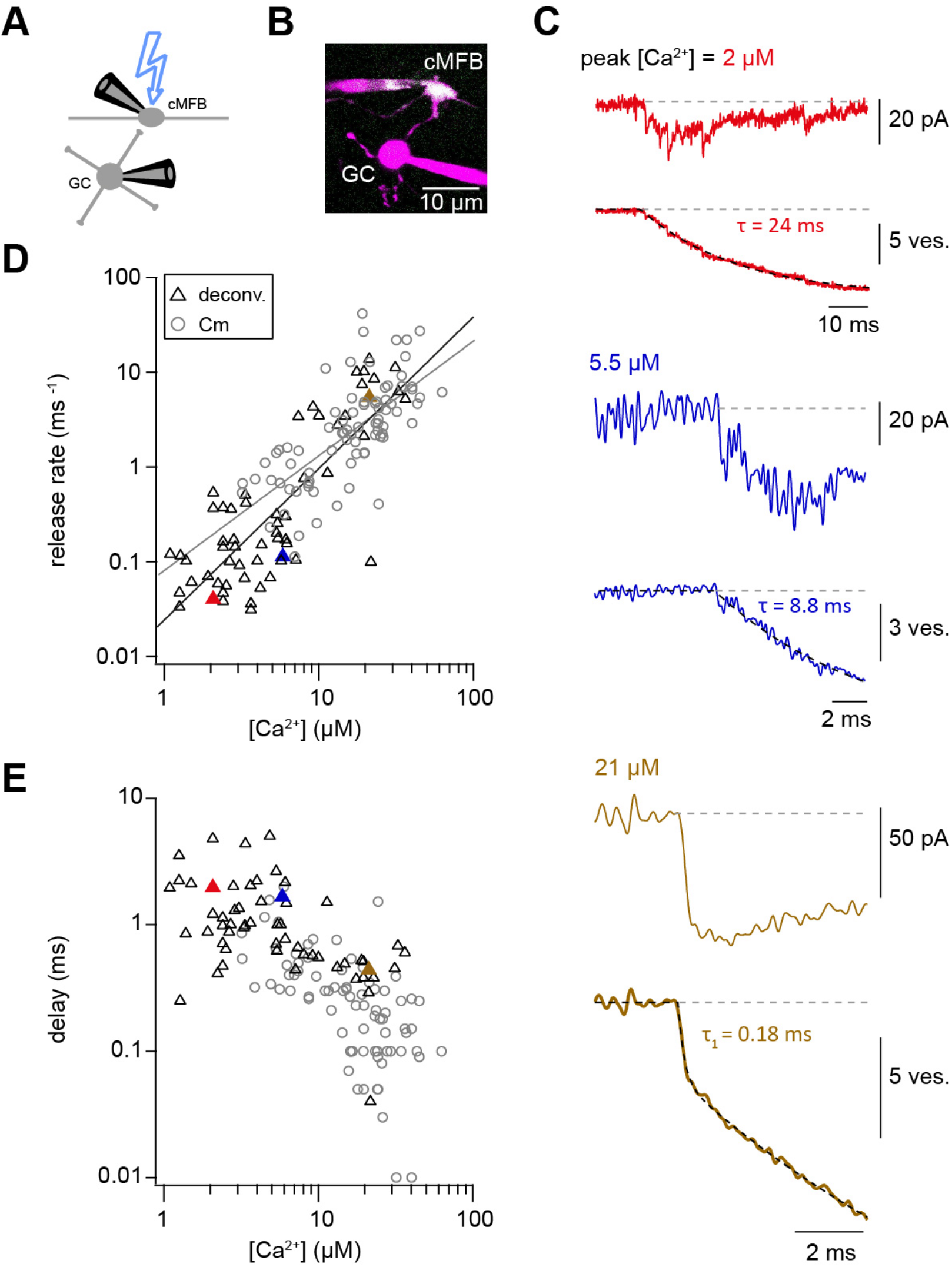
Ca^2+^ uncaging dose-response curve measured with deconvolution of EPSCs. A. Illustration of the cellular connectivity in the cerebellar cortex showing the pre- and postsynaptic compartments during paired whole-cell patch-clamp recordings and Ca^2+^ uncaging with UV-illumination. B. Two-photon microscopic image of a cMFB and a GC in the paired whole-cell patch-clamp configuration C. Three different recordings showing UV-flash evoked EPSC (*top trace*) and cumulative release rate measured by deconvolution analysis of the EPSCs (*bottom trace*). The peak Ca^2+^ concentration, quantified with two-photon Ca^2+^ imaging, is indicated in each panel. *τ* represents the time constant from mono-exponential fit, *τ*_1_ represents the time constant of the fast component of bi-exponential fit. Note the different lengths of the baselines in the three recordings. D. Plot of release rate versus post-flash Ca^2+^ concentration. Grey open circles represent data from capacitance measurements (cf. Fig. 2) and black triangles represent data from deconvolution analysis of EPSC. Grey and black lines represent fits with a Hill equation of the capacitance (as shown in Fig. 1F) and the deconvolution data, respectively. The best-fit parameters for the fit on the deconvolution data were *V_max_* = 6*10^7^ ms^-1^, *K_D_* = 7.6*10^5^ μM, and *n* = 1.6. Red, blue and brown symbols correspond to the traces in (C). E. Plot of synaptic delay versus post-flash Ca^2+^ concentration. Grey open circles represent data from capacitance measurements, and black triangles represent data from deconvolution analysis of EPSC. Red, blue and brown symbols correspond to the traces in (C).

**Figure 4.**
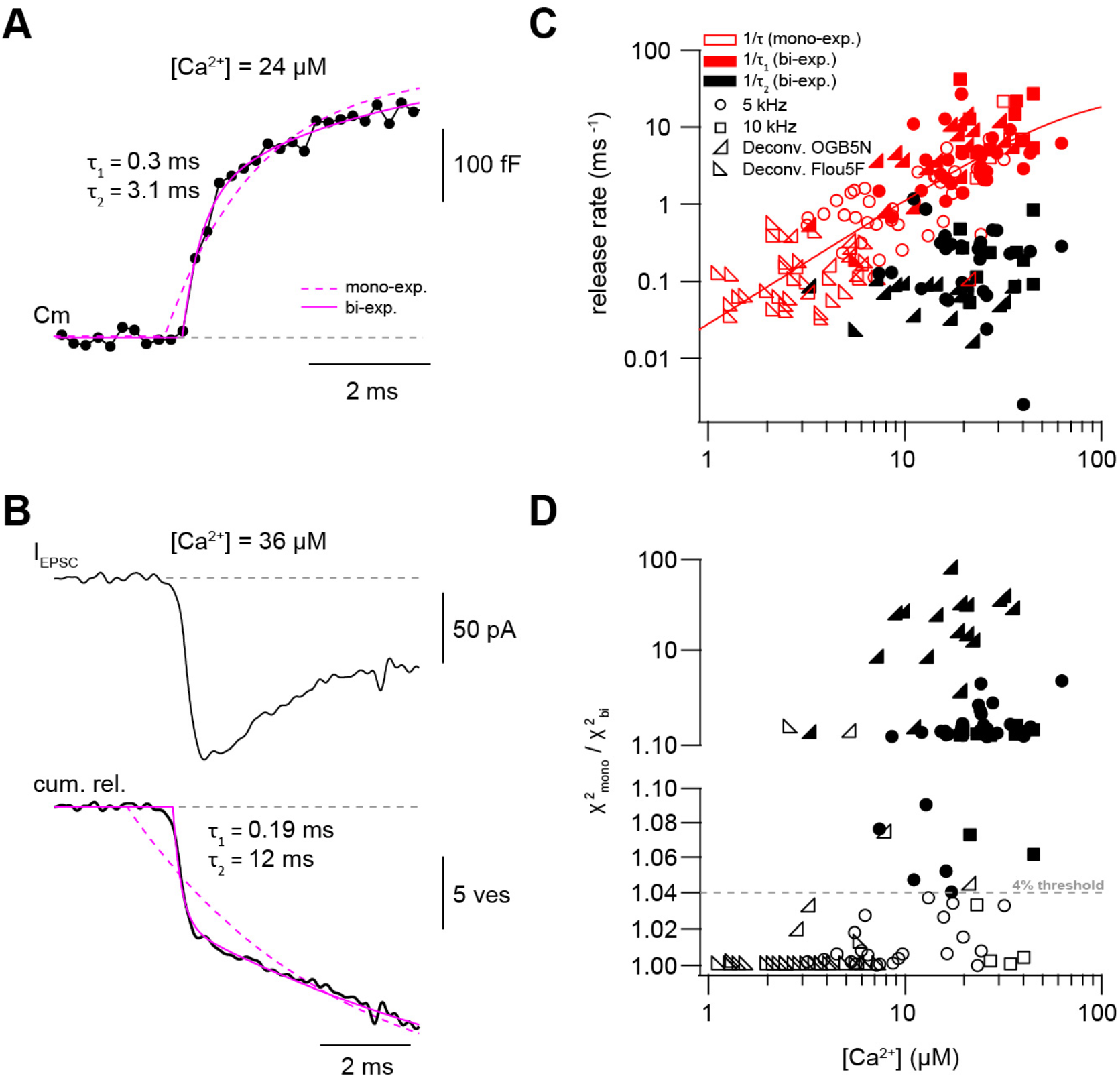
Presynaptic and postsynaptic measurements reveal two kinetic processes of neurotransmitter release. A. Example of a capacitance trace showing the two components of release observed within the first 10 ms in response to UV-flash-evoked increase in Ca^2+^ concentration to 24 μM. The solid magenta line represents the bi-exponential fit and the dashed magenta line represents mono-exponential fit (see eq. 1). B*. Top:* example trace of an EPSC recording in response to UV-flash evoked increase in Ca^2+^ concentration to 36 μM. *Bottom:* the corresponding cumulative release trace obtained from deconvolution analysis, showing the two components of release observed within the first 10 ms. The solid magenta line represents the bi-exponential fit and the dashed magenta line represents mono-exponential fit (see eq. 1). *C. Top:* plot of neurotransmitter release rates as a function of peak Ca^2+^ concentration. Data obtained from capacitance measurements with sinusoidal frequency of 5 kHz are shown as circles, data from 10 kHz capacitance measurements are shown as squares, and cumulative release data (obtained from deconvolution analysis) are shown as lower left- and lower right-triangles for recordings with OGB5N and Fluo5F, respectively. Open symbols correspond to data from the mono-exponential fits and filled symbols correspond to data from the bi-exponential fits. Red symbols represent merged data of the release rates obtained from mono-exponential fit and the fast component of the bi-exponential fit, and black symbols represent the second component of the bi-exponential fit. The line represents a fit with a Hill equation with best-fit parameters *V_max_* = 29.9 ms^-1^, *K_D_* = 75.5 μM, and *n* = 1.61. D. χ^2^ ratio for the mono-exponential compared to the bi-exponential fits. Dashed line represents the threshold of the χ^2^ ratio used to judge the fit quality of double compared to mono-exponential fits (as one criterion for selection). 5 kHz capacitance data are shown as circles, 10 kHz capacitance data are shown as squares, and cumulative release data (obtained from deconvolution analysis) are shown as lower left- and lower right-triangles for recordings with OGB5N and Fluo5F, respectively. Open symbols correspond to data points judged as mono-exponential and filled symbols correspond to data points judged as bi-exponential.

**Figure 5.**
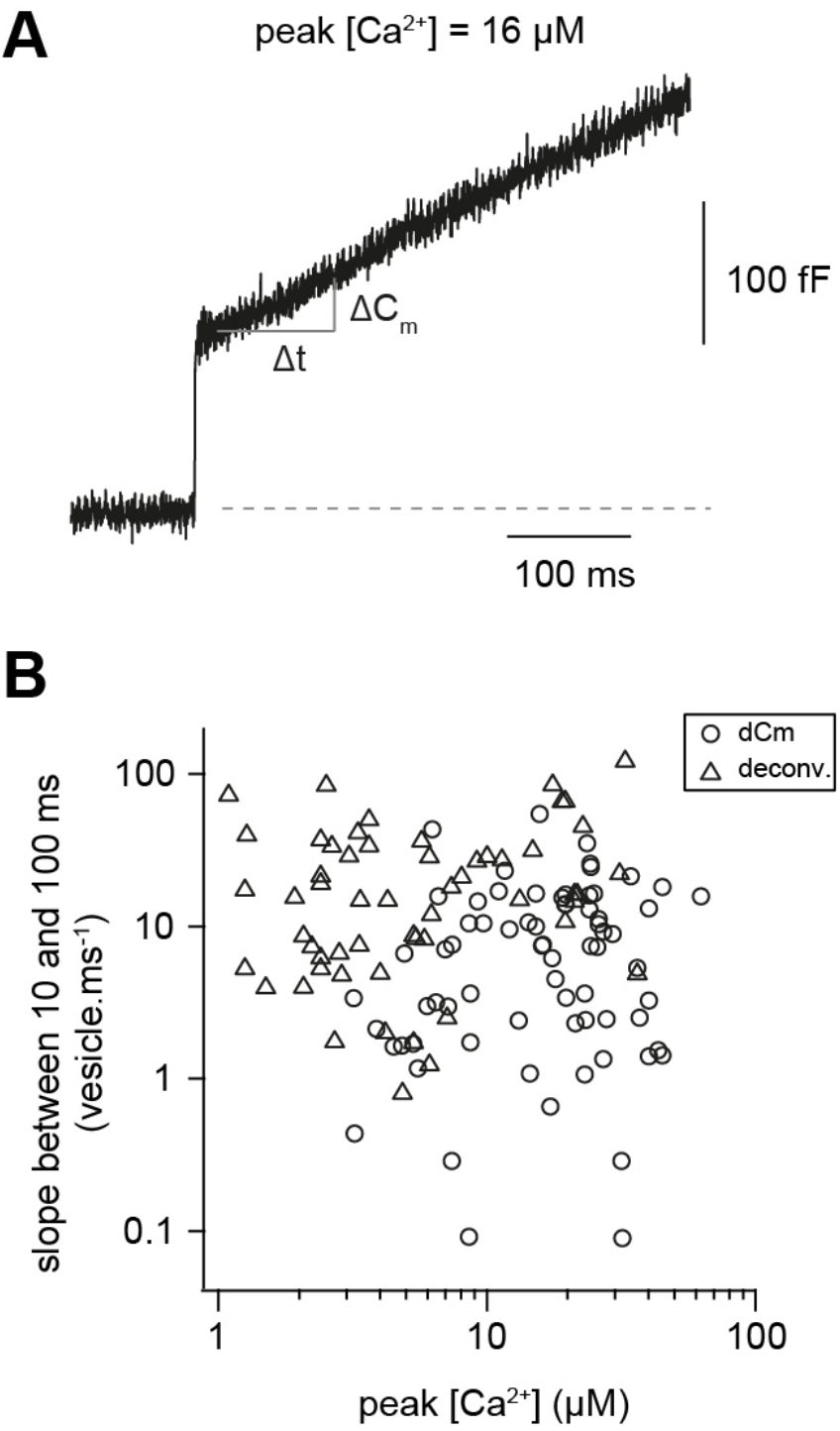
Fast and Ca^2+^-independent sustained release. A. Examples of capacitance traces showing the sustained component of release. B. Plot of the number of vesicles released between 10 and 100 ms divided by the time interval (90 ms) versus the post-flash Ca^2+^ concentration. Open circles represent data from capacitance measurements and triangles represent cumulative release data (obtained from deconvolution analysis).

**Figure 6.**
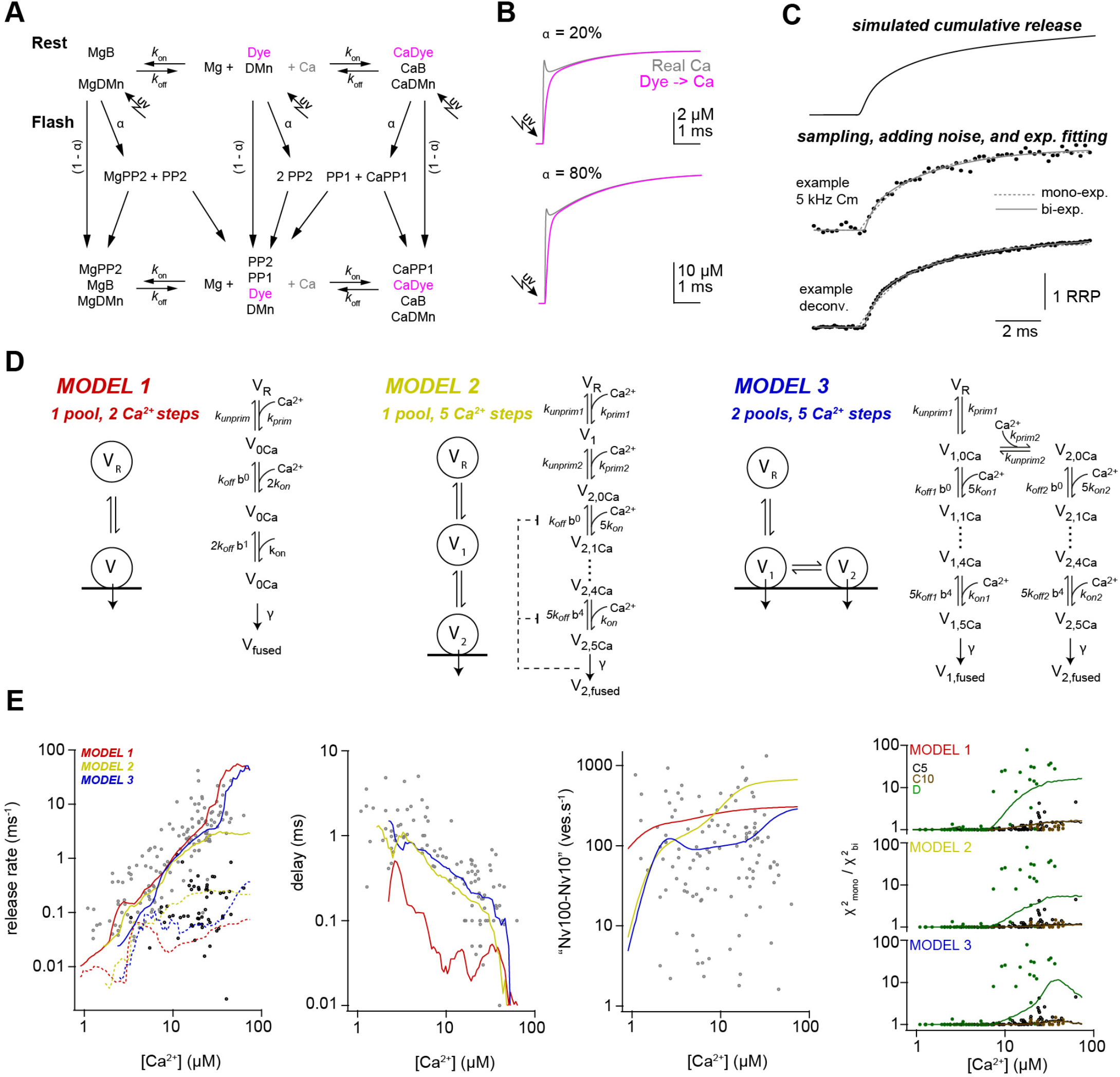
Release schemes with five Ca^2+^ steps and fast recruitment via parallel or sequential models can explain Ca^2+^-dependence of release. A. Scheme of the modeling of the intra-bouton Ca^2+^ dynamics showing the chemical reaction kinetics that were implemented in the model. The model covered Ca^2+^ (Ca) and Mg^2+^ (Mg) binding to the indicator dye (OGB-5N or Fluo-5F), to DM-nitrophen (DMn), and to buffers (ATP and/or an endogenous buffer). The forward (*K*_on_) and backward (*k*_off_) rate constants differ between chemical species. Upon simulated UV flash photolysis, a fraction α of metal bound and free DMn made a transition to different photoproducts (PP1 and PP2; cf. Faas et al., 2005). For model parameters see Supplementary Table 2. B. The scheme in (A) was converted to a system of differential equations and the time courses of the “real” free Ca^2+^ (magenta) and the free Ca^2+^ reported by OGB-5N (200 μM, green) were simulated for the indicated uncaging fractions a. Note that already after less than 1 ms the dye reliably reflects the time course of Ca^2+^. C. Traces showing the steps used in the simulation of the kinetic model of release. D. Graphical illustration of the three models used during the simulations. For model parameters see Supplementary Table 3. E. From left to right, predictions of each model and the experimental data for the inverse of τ_1_ (grey symbols, solid lines) and inverse of τ_2_ (black symbols, dashed lines), delay, vesicle recruitment speed between 10 and 100 ms, and the increase in the χ^2^ ratio for the single-compared to the bi-exponential fits. Red, yellow, and blue lines correspond to simulations of models 1,2, and 3, respectively. For the χ^2^ ratio (*right plot*), the experimental data and the simulations are shown separately for 5-kHz and 10-kHz capacitance data (C5 and C10; black and brown, respectively) and the deconvolution data (D; green).

The suffixes “T”, T, and “s” indicate total, fast or slow, respectively. The transition of fast and slow uncaging fractions into low-affinity photoproducts (PP) occurred with fast (τ_f_) or slow (τ_s_) time constants, respectively. Free Ca^2+^ or Mg^2+^-bound DMn decomposed into two photoproducts (PP1, PP2) differing with respect to their binding kinetics. The binding kinetics of all species were governed by the corresponding forward (*K_on_*) and backward (*K_off_*) rate constants

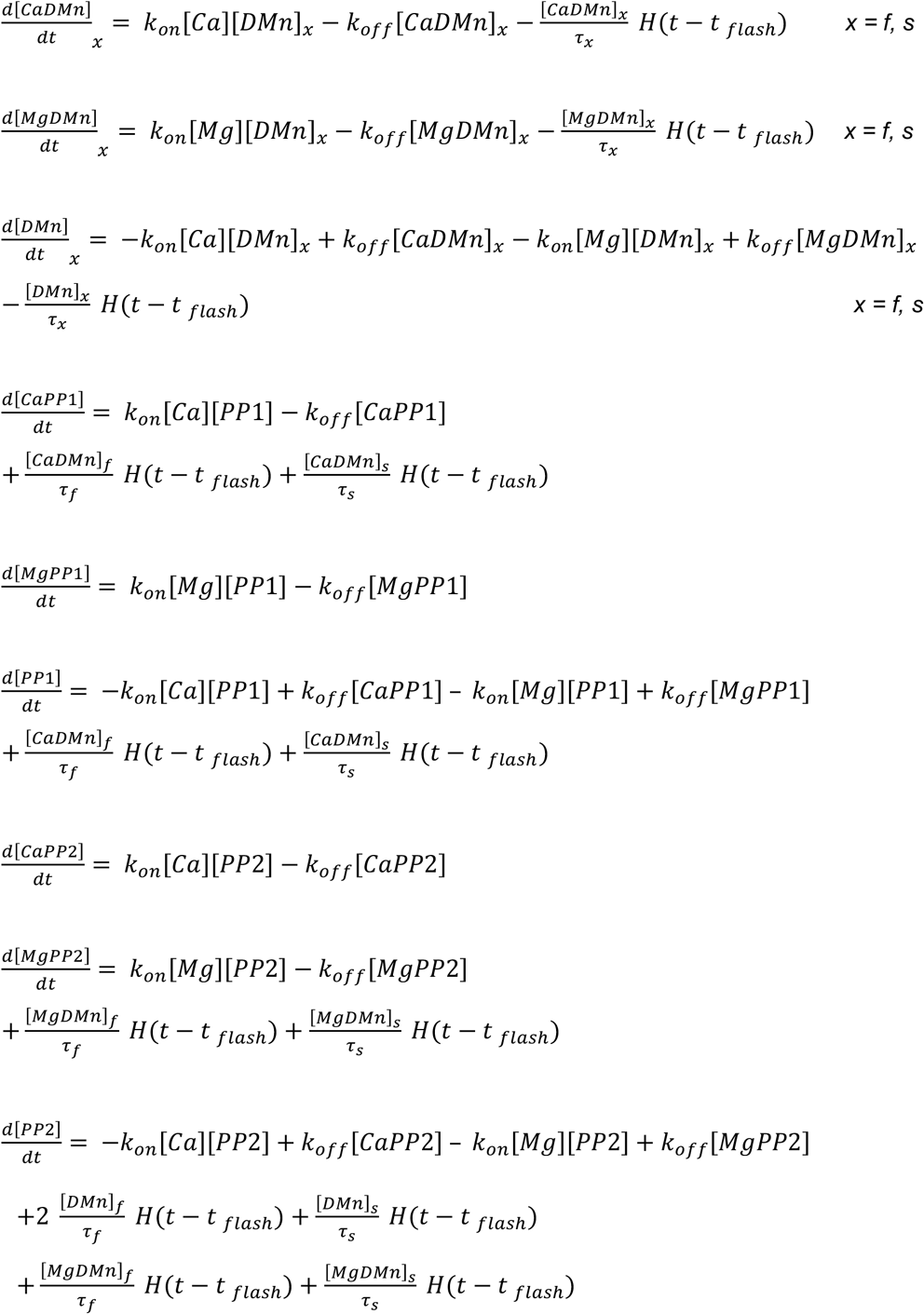

where *H* is the Heaviside step function and *t_flash_* the time of the UV flash. Ca^2+^ and Mg^2+^ binding to the dye, ATP, and an endogenous buffer (EB) were simulated by second order kinetics:

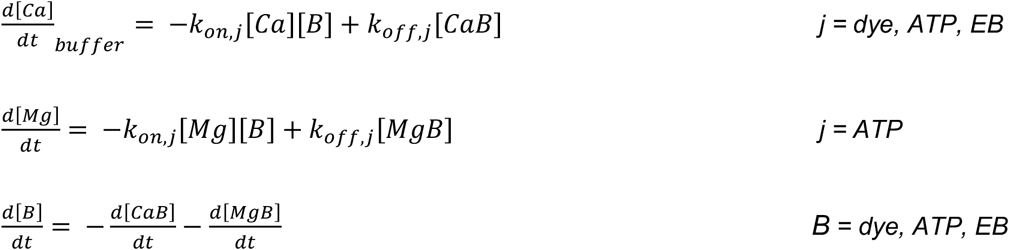

The time course of the total change Ca^2+^ concentration or Mg^2+^ concentration is given by the sum of all the above equations involving changes in Ca^2+^ concentration or Mg^2+^ concentration, respectively. Ca^2+^ concentration as reported by the dye was calculated from the concentration of the Ca^2+^-dye complex assuming equilibrium conditions (Markram et al., 1998). The clearing of Ca^2+^ from the cytosol was not implemented in these simulations. Instead, the Ca^2+^ concentration was simulated only for 10 ms after the flash. The experimentally observed subsequent decay of the Ca^2+^ concentration was implemented by an exponential decay to the resting Ca^2+^ concentration with a time constant of 400 ms. The parameters of the model are given in Supplementary Table 2.

**Supplementary Table 2.** Parameters for simulations of Ca^2+^ release from DMN cage.

| Parameters |  | Values | References number / Notes |
| --- | --- | --- | --- |
| Resting $Ca^{2+}$ | $[Ca^{2+}]_{rest}$ | $227 \cdot 10^{-9}$ M | Measured |
| Total magnesium | $[Mg^{2+}]_T$ | $0.5 \cdot 10^{-3}$ M | Pipette concentration |
| Fluo-5F | $[Fluo]$ | 0 or $50 \cdot 10^{-6}$ M (see Supplementary Table 1) | Pipette concentration |
| | $K_D$ | $0.83 \cdot 10^{-6}$ M | (Delvendahl et al., 2015) |
| | $k_{\text{off}}$ | 249 s<sup>-1</sup> | ibid |
| | $k_{\text{on}}$ | $3 \cdot 10^8 \text{ M}^{-1} \text{ s}^{-1}$ | (Yasuda et al., 2004) |
| OGB-5N | [OGB] | 0 or $200 \cdot 10^{-6} \text{ M}$ (see Supplementary Table 1) | Pipette concentration |
| | $K_D$ | $24 \cdot 10^{-6} \text{ M}$ | (Delvendahl et al., 2015) |
| | $k_{\text{off}}$ | 6000 s<sup>-1</sup> | ibid. |
| | $k_{\text{on}}$ | $2.5 \cdot 10^8 \text{ M}^{-1} \text{ s}^{-1}$ | (Digregorio and Vergara, 1997) |
| ATP | [ATP] | $5 \cdot 10^{-3} \text{ M}$ | Pipette concentration |
| Ca<sup>2+</sup> binding | $K_D$ | $2 \cdot 10^{-4} \text{ M}$ | (Meinrenken et al., 2002) |
| | $k_{\text{off}}$ | 100 000 s<sup>-1</sup> | ibid. |
| | $k_{\text{on}}$ | $5 \cdot 10^8 \text{ M}^{-1} \text{ s}^{-1}$ | ibid. |
| Mg<sup>2+</sup> binding | $K_D$ | $100 \cdot 10^{-6} \text{ M}$ | (Bollmann et al., 2000); MaxC |
| | $k_{\text{off}}$ | 1000 s<sup>-1</sup> | ibid. |
| | $k_{\text{on}}$ | $1 \cdot 10^7 \text{ M}^{-1} \text{ s}^{-1}$ | ibid. |
| Endogenous buffer | [EB] | $480 \cdot 10^{-6} \text{ M}$ | (Delvendahl et al., 2015) |
| | $K_D$ | $32 \cdot 10^{-6} \text{ M}$ | ibid |
| | $k_{\text{off}}$ | 16 000 s<sup>-1</sup> | ibid. |
| | $k_{\text{on}}$ | $5 \cdot 10^8 \text{ M}^{-1} \text{ s}^{-1}$ | ibid. |
| Total DM nitrophen | [DMn]<sub>T</sub> | $500 \cdot 10^{-6} - 10 \cdot 10^{-3} \text{ M}$ (see Supplementary Table 1) | Pipette concentration |
| Ca<sup>2+</sup> binding | $K_D$ | $6.5 \cdot 10^{-9} \text{ M}$ | (Faas et al., 2005) |
| | $k_{\text{off}}$ | 0.19 s<sup>-1</sup> | ibid. |
| | $k_{\text{on}}$ | $2.9 \cdot 10^7 \text{ M}^{-1} \text{ s}^{-1}$ | ibid. |
| Mg<sup>2+</sup> binding | $K_D$ | $1.5 \cdot 10^{-6} \text{ M}$ | ibid. |
| | $k_{\text{off}}$ | 0.2 s<sup>-1</sup> | ibid. |
| Uncaging fraction | $\alpha$ | See Supplementary Table 1 | |
| Fast uncaging fraction | af | 0.67 | (Faas et al., 2005) |
| Photoproduct 1 | [PP1] |  |  |
| Ca<sup>2+</sup> binding | $K_D$ | $2.38 \cdot 10^{-3} \text{ M}$ | (Faas et al., 2005) |
| | $k_{\text{off}}$ | 69 000 s<sup>-1</sup> | ibid. |
| | $k_{\text{on}}$ | $2.9 \cdot 10^7 \text{ M}^{-1} \text{ s}^{-1}$ | ibid. |
| Mg<sup>2+</sup> binding | $K_D$ | $1.5 \cdot 10^{-6} \text{ M}$ | ibid. |
| | $k_{\text{off}}$ | 300 s<sup>-1</sup> | ibid. |
| | $k_{\text{on}}$ | $1.3 \cdot 10^5 \text{ M}^{-1} \text{ s}^{-1}$ | ibid. |
| Photoproduct 2 | [PP2] |  |  |
| Ca<sup>2+</sup> binding | $K_D$ | $124.1 \cdot 10^{-6} \text{ M}$ | ibid. |
| | $k_{off}$ | $3600 \text{ s}^{-1}$ | ibid. |
| | $k_{on}$ | $2.9 \cdot 10^7 \text{ M}^{-1} \text{ s}^{-1}$ | ibid. |
| Mg<sup>2+</sup> binding | $K_D$ | $1.5 \cdot 10^{-6} \text{ M}$ | ibid. |
| | $k_{off}$ | $300 \text{ s}^{-1}$ | ibid. |
| | $k_{on}$ | $1.3 \cdot 10^5 \text{ M}^{-1} \text{ s}^{-1}$ | ibid. |

These simulations were used to obtain Ca^2+^ transients with peak amplitudes covering the entire range of post-flash Ca^2+^ concentrations. To this end, the uncaging efficiency a was varied in each of the three experimentally used combinations of concentrations of DMn and Ca^2+^ indicators (see Supplementary Table 1 for details).

### Modeling of release schemes

Model 1 with two Ca^2+^ binding steps mediating fusion and one Ca^2+^-dependent priming step was defined according to the following differential equation

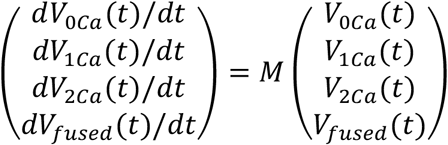

*V_0Ca_*, *V*_1*Ca*_, and *V*_2*Ca*_ denote the fraction of vesicles with a fusion sensor with 0 to 2 bound Ca^2+^ ions, respectively, and *V_fused_* denotes the fused vesicles as illustrated in Fig. 6D. The reserve pool *V_R_* is considered to be infinite. *M* denotes the following 4×4 matrix:

|  |  |  |  |
| --- | --- | --- | --- |
| $-2k_{on} - k_{unprim} + k_{prim}/V_{0Ca}(t)$ | $k_{off}$ | 0 | 0 |
| $2k_{on}$ | $-k_{off} - k_{on}$ | $2k_{off}b$ | 0 |
| 0 | $k_{on}$ | $-\gamma - 2k_{off}b$ | 0 |
| 0 | 0 | $\gamma$ | 0 |

See Supplementary Table 3 for the values and Ca^2+^-dependence of the rate constants in the matrix.

The initial condition was defined as *V*_0*Ca*_(0) = *K_prim_*/*K_unprim_* and *V*_1*Ca*_(0), *V*_2*Ca*_(0), and *V_fused_*(0) was zero. *K_prim_* was the sum of a Ca^2+^-dependent and Ca^2+^-independent rate constants. The Ca^2+^-dependence was implemented as a Michaelis-Menten kinetic with a maximum rate constant of 30 s^-1^ (Ritzau-Jost et al., 2014) and a *K_D_* of 2 μM (Miki et al., 2018). The Ca^2+^-independent rate constant was 0.6 s^-1^, adjusted to reproduce the factor of 3 upon elevating Ca^2+^ from 30 to 180 nM (cf. Fig. 1D and 7D). *K*_unprim_ was defined such that the occupancy *V*_0*Ca*_ (0) = 1 for the default pre-flash resting Ca^2+^ concentration of 227 nM (Supplementary Tables 2 and 3).

**Figure 7.**
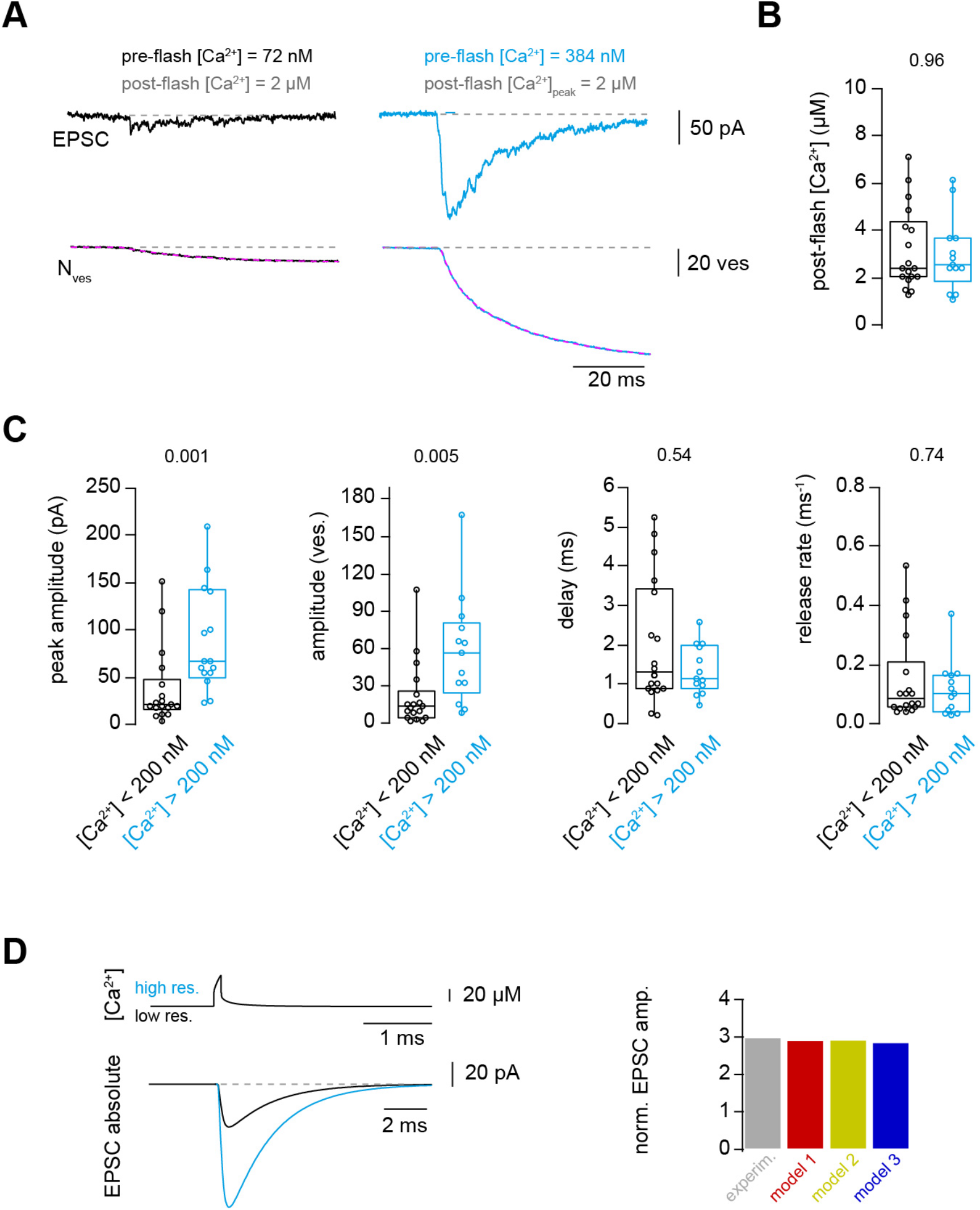
Ca^2+^ uncaging with different pre-flash Ca^2+^ concentrations indicates Ca^2+^-dependent vesicle priming. A. Two consecutive recordings from the same cell pair, with the same post-flash Ca^2+^ concentration but different pre-flash Ca^2+^ concentration in the presynaptic terminal. *Top:* postsynaptic current. *Bottom:* cumulative release of synaptic vesicles measured by deconvolution analysis of EPSCs superposed with a mono-exponential fit (magenta). Black and blue color represent low and high pre-flash Ca^2+^ concentration, respectively. The pre- and post-flash Ca^2+^ concentrations are indicated in each panel. B. Comparison of the average post-flash Ca^2+^ concentration between both groups of either low or high pre-flash Ca^2+^ concentration (black and blue bars, respectively). C. From left to right: comparisons of the peak amplitude, the number of released vesicles measured as obtained from deconvolution analysis of EPSC, the delay of the release onset, and the release rate. Boxplots show median and 1^st^/3^rd^ quartiles with whiskers indicating the whole data range. The values above the boxplots represent P-values of Mann-Whitney U test. D. *Top left:* simulated local Ca^2+^ signal at 20 nm from the Ca^2+^ channel taken from Delvendahl et al., 2015. Note the almost complete overlap of the two Ca^2+^ concertation traces with low and high basal pre-flash Ca^2+^ concertation. *Bottom left:* prediction of the increase in the amplitude of action potential-evoked EPSC, upon elevating the basal Ca^2+^ concentration in the presynaptic terminal. *Right:* comparison between experimental data and the models’ predictions of the effect of basal Ca^2+^ on the amplitude of the action potential-evoked release.

The differential equations were solved with the NDSolve function of Mathematica. The Ca^2+^ concentration, Ca^2+^(*t*), was obtained from the simulations as described in the previous paragraph. *V_fused_*(*t*) represents the cumulative release normalized to the pool of release-ready vesicles per cMFB to GC connection. To reproduce the absolute sustained release rate (Figs. 5 and 6D), *V_fused_*(*t*) was multiplied by a pool of release-ready vesicles per connection of 10 vesicles. The cumulative release, *V_fused_*(*t*), including a pre-flash baseline was sampled with 5 or 10 kHz. Realistic noise for 5- or 10 kHz-capacitance or deconvolution measurements was added and the data, in the 10 ms-window after the flash, were fit with mono- and bi-exponential functions (eq. 1). The selection of a bi-over a mono-exponential fit was based on identical criteria as in the analysis of the experimental data including the prolongation of the fitting duration from 10 to 60 ms if the time constant of the mono-exponential fit was >10 ms (see section Data analysis). For each peak post-flash Ca^2+^ concentration (i.e. simulated Ca^2+^(*t*) transient) the sampling, addition of noise, and exponential fitting were repeated 50 times. The median of these values represents the prediction of the model for each peak post flash Ca^2+^ concentration. The parameters of the models were manually adjusted to obtain best-fit results.

Model 2 was a sequential two-pool model based on Miki et al. (2018) with five Ca^2+^ binding steps mediating fusion and two Ca^2+^-dependent priming steps defined according to the following differential equations

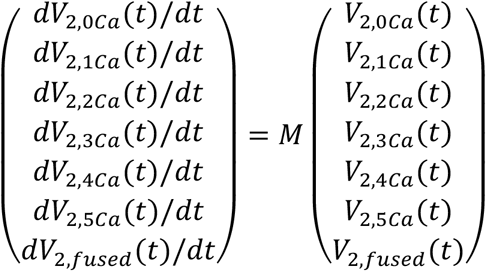

*V*_2,0*Ca*_, *V*_2,1*Ca*_,…, and *V*_2,5*Ca*_ denote the fraction of vesicles with a fusion sensor with 0 to 5 bound Ca^2+^ ions, respectively, and *V*_2,*fused*_ denotes fused vesicles as illustrated in Fig. 6D. The fraction of vesicles in state *V*_1_ is calculated according to the following differential equation

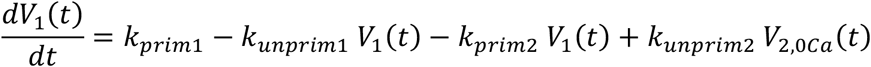

*M* denotes the following 7×7 matrix:

|  |  |  |  |  |  |  |
| --- | --- | --- | --- | --- | --- | --- |
| $-5k_{on} - k_{unprim2} + k_{prim2} V_1(t)/V_{2,0Ca}(t)$ | $k_{off}$ | 0 | 0 | 0 | 0 | 0 |
| $5k_{on}$ | $-k_{off} - 4k_{on}$ | $2k_{off}b$ | 0 | 0 | 0 | 0 |
| 0 | $4k_{on}$ | $-2k_{off}b - 3k_{on}$ | $3k_{off}b^2$ | 0 | 0 | 0 |
| 0 | 0 | $3k_{on}$ | $-3k_{off}b^2 - 2k_{on}$ | $4k_{off}b^3$ | 0 | 0 |
| 0 | 0 | 0 | $2k_{on}$ | $-4k_{off}b^3 - k_{on}$ | $5k_{off}b^4$ | 0 |
| 0 | 0 | 0 | 0 | $k_{on}$ | $-\gamma - 5k_{off}b^4$ | 0 |
| 0 | 0 | 0 | 0 | 0 | $\gamma$ | 0 |

To implement the use-dependent slowing of the release rate constants of this model (Miki et al., 2018) in a deterministic way, a site-plugging state, *P*(*t*), was defined according to

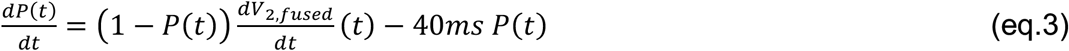

*P*(*t*) is approaching 1 during strong release and decays with a time constant of 40 ms back to zero. Similar to the implementation by Miki et al. (2018), the rate constants *K_on_* and *K_off_* were linearly interpolated between two values depending on *P*(*t*) as

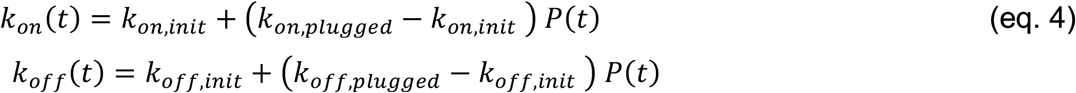

The reserve pool *V_R_* is considered to be infinite. See Supplementary Table 3 for the values and Ca^2+^-dependence of the rate constants in these differential equations.

The initial condition is defined as *V*_1_(0) = *K_prim1_*/*K_unprim1_* and *V*_2,0*Ca*_(0) = (*K_prim1_*/*K*_unprim1_)*(*K_prim2_*/*K_unprim2_*). The initial condition of the other state *V*_2,1*Ca*_(0) to *V*_5,0*Ca*_(0), *V_fused_*(0), and *P*(0) were zero. *K_prim1_* and *K_prim2_* were the sum of a Ca^2+^-dependent and Ca^2+^-independent rate constant defined similarly as described in Miki et al. (2018) and adjusted as described for model 1. *K_unprim1_* and *K_unprim2_* were defined such that the occupancy *V*_1_(0) = 1 and *V*_2,0*Ca*_(0) = 1 for the default pre-flash resting Ca^2+^ concentration of 227 nM (Supplementary Tables 2 and 3).

Model 3 was a parallel two-pool model similar as described by Voets (2000) and Walter et al (2013) but with five Ca^2+^ binding steps mediating fusion of both types of vesicles and a Ca^2+^-independent priming step for V_1_ vesicles and a Ca^2+^-dependent transition step from V_1_ to V_2_ vesicles defined according to the following differential equations

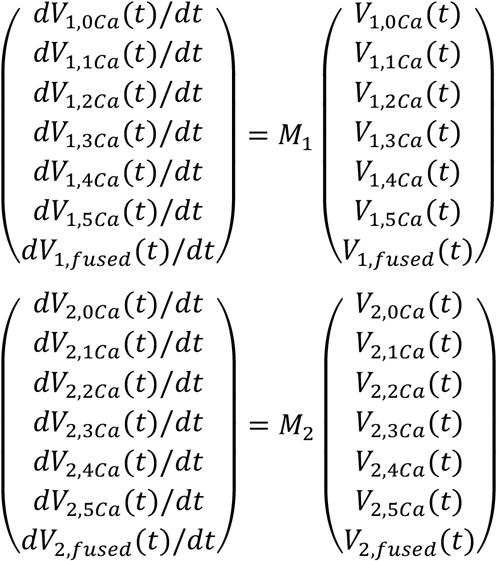

*V*_1,0*Ca*_, *V*_1,1*Ca*_,…, and *V_1,5*Ca*_* denote the fraction of vesicles with a low-affinity fusion sensor with 0 to 5 bound Ca^2+^ ions, respectively, and *V*_2,0*Ca*_, *V*_2,1*Ca*_, and *V*_2,5*Ca*_ denote the fraction of vesicles with a high-affinity fusion sensor with 0 to 5 bound Ca^2+^ ions, respectively. *V*_1,*fused*_ and *V*_2,*fused*_ denote fused vesicles as illustrated in Fig. 6D.

*M*_1_ denotes the following 7×7 matrix:

|  |  |  |  |  |  |  |
| --- | --- | --- | --- | --- | --- | --- |
| $-5k_{on1}-k_{unprim1}-k_{prim2}$<br>$+k_{prim1}/V_{1,0Ca}(t)$<br>$+k_{unprim2}V_{2,0Ca}(t)/$<br>$V_{1,0Ca}(t)$ | $k_{off1}$ | 0 | 0 | 0 | 0 | 0 |
| $5k_{on1}$ | $-k_{off1}-4k_{on1}$ | $2k_{off1}b$ | 0 | 0 | 0 | 0 |
| 0 | $4k_{on1}$ | $-2k_{off1}b-3k_{on1}$ | $3k_{off1}b^2$ | 0 | 0 | 0 |
| 0 | 0 | $3k_{on1}$ | $-3k_{off1}b^2-2k_{on1}$ | $4k_{off1}b^3$ | 0 | 0 |
| 0 | 0 | 0 | $2k_{on1}$ | $-4k_{off1}b^3-k_{on1}$ | $5k_{off1}b^4$ | 0 |
| 0 | 0 | 0 | 0 | $k_{on1}$ | $-\gamma-5k_{off1}b^4$ | 0 |
| 0 | 0 | 0 | 0 | 0 | $\gamma$ | 0 |

*M*_2_ denotes the following 7×7 matrix:

$M_2$ denotes the following 7x7 matrix:
|  |  |  |  |  |  |  |
| --- | --- | --- | --- | --- | --- | --- |
| $-5k_{on1}-k_{unprim2}$<br>$+k_{prim2}V_{1,0Ca}(t)/$<br>$V_{2,0Ca}(t)$ | $k_{off2}$ | 0 | 0 | 0 | 0 | 0 |
| $5k_{on2}$ | $-k_{off2}-4k_{on2}$ | $2k_{off2}b$ | 0 | 0 | 0 | 0 |
| 0 | $4k_{on2}$ | $-2k_{off2}b-3k_{on2}$ | $3k_{off2}b^2$ | 0 | 0 | 0 |
| 0 | 0 | $3k_{on2}$ | $-3k_{off2}b^2-2k_{on2}$ | $4k_{off2}b^3$ | 0 | 0 |
| 0 | 0 | 0 | $2k_{on2}$ | $-4k_{off2}b^3-k_{on2}$ | $5k_{off2}b^4$ | 0 |
| 0 | 0 | 0 | 0 | $k_{on2}$ | $-\gamma-5k_{off2}b^4$ | 0 |
| 0 | 0 | 0 | 0 | 0 | $\gamma$ | 0 |

The initial condition is defined as *V*_2,0*Ca*_(0) = *K_prim1_*/*K_unprim1_* and *V*_2,0*Ca*_(0) = (*K_prim1_*/*K*_unprim1_)*(*K*_prim2_/*K_unprim2_*). The initial condition of the other state *V*_1,1*Ca*_(0) to *V*_1,0*Ca*_(0), *V*_1,*fused*_(0), and *V*_2,1*Ca*_(0) to *V*_2,0*Ca*_(0), *V*_2,*fused*_(0) were zero, *K_prim1_* was a Ca^2+^-independent rate constant and *K_prim2_* was the sum of a Ca^2+^-dependent and Ca^2+^-independent rate constants defined similarly as described in Hallermann et al. (2010) and adjusted as described for model 1. *K_unprim1_* and *K_unprim2_* were defined such that the occupancy *V*_1,0*Ca*_(0) = 1 and *V*_2,0*Ca*_(0) = 1 for the default pre-flash resting Ca^2+^ concentration of 227 nM (Supplementary Tables 2 and 3).

**Supplementary Table 3.** Parameters for release scheme models.

| Model1 |  | Model2 |  | Model3 |  |
| --- | --- | --- | --- | --- | --- |
| $k_{on}$ | $2.95 \cdot 10^9 Ca^{2+}(t) M^{-1} s^{-1}$ | $k_{on,init}$ | $5.10 \cdot 10^8 Ca^{2+}(t) M^{-1} s^{-1}$ | $k_{on1}$ | $0.5 k_{on2}$ |
| | | $k_{on,plug}$ | $0.1 k_{on,init}$ | $k_{on2}$ | $5.10 \cdot 10^8 Ca^{2+}(t) M^{-1} s^{-1}$ |
| $k_{off}$ | $4.42 \cdot 10^5 s^{-1}$ | $k_{off,init}$ | $2.55 \cdot 10^4 s^{-1}$ | $k_{off1}$ | $10 k_{off2}$ |
| | | $k_{off,plug}$ | $0.4 k_{off,init}$ | $k_{off2}$ | $2.55 \cdot 10^4 s^{-1}$ |
| b | 0.25 | b | 0.25 | b | 0.25 |
| $\gamma$ | $1.77 \cdot 10^4 s^{-1}$ | $\gamma$ | $1.77 \cdot 10^4 s^{-1}$ | $\gamma$ | $1.77 \cdot 10^4 s^{-1}$ |
| $k_{prim}$ | $0.6 + 30 \cdot (Ca^{2+}(t) / (K_{d,prim} + Ca^{2+}(t))) s^{-1}$ | $k_{prim1}$ | $2.5 + 60 \cdot (Ca^{2+}(t) / (K_{d,prim1} + Ca^{2+}(t))) s^{-1}$ | $k_{prim1}$ | $30 s^{-1}$ |
| $k_{unprim}$ | $0.6 + 30 \cdot (Ca^{2+}_{Rest} / (K_{d,prim} + Ca^{2+}_{Rest})) s^{-1}$ | $k_{unprim1}$ | $2.5 + 60 \cdot (Ca^{2+}_{Rest} / (K_{d,prim1} + Ca^{2+}_{Rest})) s^{-1}$ | $k_{unprim1}$ | $30 s^{-1}$ |
| $K_{d,prim}$ | 2 $\mu M$ | $K_{d,prim1}$ | 2 $\mu M$ | | |
| | | $k_{prim2}$ | $100 + 800 \cdot (Ca^{2+}(t) / (K_{d,prim2} + Ca^{2+}(t))) s^{-1}$ | $k_{prim2}$ | $0.5 + 30 \cdot (Ca^{2+}(t) / (K_{d,prim2} + Ca^{2+}(t))) s^{-1}$ |
| | | $k_{unprim2}$ | $100 + 800 \cdot (Ca^{2+}_{Rest} / (K_{d,prim2} + Ca^{2+}_{Rest})) s^{-1}$ | $k_{unprim2}$ | $0.5 + 30 \cdot (Ca^{2+}_{Rest} / (K_{d,prim2} + Ca^{2+}_{Rest})) s^{-1}$ |
| | | $K_{d,prim2}$ | 2 $\mu M$ | $K_{d,prim2}$ | 2 $\mu M$ |

### Statistical analysis

Boxplots show median and 1st/3rd quartiles with whiskers indicating the whole data range (Figs. 1 and 7). For statistical comparison, Mann-Whitney U tests were used, and the P-values are indicated above the boxplots.

## Results

### Action potential-evoked synaptic release critically depends on basal intracellular Ca^2+^ concentration

To investigate the impact of the basal intracellular Ca^2+^ concentration on synaptic release, we performed simultaneous patch-clamp recordings from presynaptic cerebellar mossy fiber boutons (cMFB) and postsynaptic granule cells (GC) of 5-to 6-weeks old mice at physiological temperatures (Fig. 1A and B). We aimed at clamping the free Ca^2+^ concentration in the presynaptic patch solution to either low or high basal Ca^2+^ concentrations by adding different concentrations of Ca^2+^ and the Ca^2+^ chelator EGTA (see methods). Two-photon quantitative Ca^2+^ imaging with the dual-indicator method using Fluo-5F as the Ca^2+^ indicator (Delvendahl et al., 2015; Sabatini et al., 2002) revealed the free Ca^2+^ concentration of the presynaptic intracellular solution to be 28 ± 3 and 183 ± 8 nM, for the low and high basal Ca^2+^ conditions (n = 4 and 4), respectively (Fig. 1A). In both solutions, the free EGTA concentration was 4.47 mM (see methods). In response to triggering a single action potential in the presynaptic terminal, the recorded excitatory postsynaptic current (EPSC) depended strongly on the presynaptic resting Ca^2+^ concentration (Fig. 1C). We found an almost three-fold increase in the EPSC amplitude when elevating the resting Ca^2+^ concentration in the presynaptic terminals from 30 to 180 nM. On average, the EPSC amplitudes were 39 ± 8 and 117 ± 27 pA for the low and high basal Ca^2+^ conditions, respectively (n = 8 and 8; P_Mann-Whitney_ = 0.028; Fig. 1D). The EPSC rise and decay kinetics were not significantly different (Fig. 1D). No significant differences were observed in the action potential waveform including amplitude and half duration (Fig. 1D) indicating that the altered synaptic strength was not caused by changes in the shape of the presynaptic action potential. These data indicate that moderate changes in the presynaptic basal Ca^2+^ concentration can alter synaptic strength up to three-fold.

Boxplots show median and 1^st^/3^rd^ quartiles with whiskers indicating the whole data range. Values of individual experiments are superimposed as circles. The numbers above the boxplots represent P-values of Mann-Whitney U tests.

### Ca^2+^ uncaging dose-response curve measured with presynaptic capacitance measurements

To gain a better understanding of the profound sensitivity of AP-evoked release on presynaptic basal Ca^2+^ concentration, we established presynaptic Ca^2+^ uncaging and measured the release kinetics upon step-wise elevation of Ca^2+^ concentration. We combined wide-field illumination using a high-power UV laser with previously established quantitative two-photon Ca^2+^ imaging (Delvendahl et al., 2015) to quantify the post-flash Ca^2+^ concentration (Fig. 2A). This approach offers sub-millisecond control of the UV flashes and a high signal to noise ratio of the two-photon Ca^2+^ imaging deep within the brain slice. The flash-evoked artefacts in the two-photon signals, presumably due to luminescence in the light path, could be reduced to a minimum with an optimal set of spectral filters and gate-able photomultipliers (PMTs). Subtraction of the remaining artefact in the background region of the two-photon line scan resulted in artefact-free fluorescence signals (Fig. 2B and C).

To obtain a large range of post-flash Ca^2+^ concentrations within the bouton, we varied the concentration of the Ca^2+^-cage DMn (1-10 mM) and the intensity (10 – 100%) and the duration (100 or 200 μs) of the UV laser pulse. The spatial homogeneity of the Ca^2+^ elevation was assessed by UV illumination of caged fluorescein mixed with glycerol (Fig. 2 – figure supplement 1; Schneggenburger et al., 2000; Bollmann et al., 2000). The resulting post-flash Ca^2+^ concentration was quantified with either high- or low-affinity Ca^2+^ indicator (Fluo-5F or OGB-5N). To measure the kinetics of neurotransmitter release independent of dendritic filtering or postsynaptic receptor saturation, vesicular fusion was quantified by measuring the presynaptic capacitance with a 5 kHz-sinusoidal stimulation (Hallermann et al., 2003). The first 10 ms of the flash-evoked capacitance increase was fitted with functions containing a baseline and mono- or bi-exponential components (magenta line in Fig. 2D and E; see eq. 1 in the methods section). With increasing post-flash Ca^2+^ concentration the fast time constant decreased (τ in case of mono- and τ1 in case of bi-exponential fits; Fig. 2D). The inverse of the fast time constant represents a direct readout of the fusion kinetics of the release-ready vesicles. The observed scatter could be due to the invasiveness of presynaptic recordings and/or heterogeneity among boutons (Chabrol et al., 2015; Fekete et al., 2019; Grande and Wang, 2011). When plotting the inverse of the time constant as a function of post-flash Ca^2+^ concentration, we obtained a shallow dose-response curve that showed a continuous increase in the release rate with increasing post-flash Ca^2+^ concentration up to 50 μM (Fig. 2F). In some experiments with high Ca^2+^ concentrations, the release was too fast to be resolved with 5 kHz capacitance sampling (i.e. time constants were smaller than 200 μs; Fig. 2E). We therefore increased the frequency of the sinusoidal stimulation in a subset of experiments to 10 kHz (15 out of 80 experiments). Such high-frequency capacitance sampling is to our knowledge unprecedented at central synapses and technically challenging because exceptionally low access resistances are required (<~15 MΩ) to obtain an acceptable signal-to-noise ratio (Gillis, 1995; Hallermann et al., 2003). Despite these efforts, the time constants were sometimes faster than 100 μs, representing the resolution limit of 10 kHz capacitance sampling (Fig. 2E). These results indicate that the entire pool of release-ready vesicles can fuse within less than 100 μs. Fitting a Hill equation on both 5- and 10 kHz data resulted in a best-fit *K_D_* of >50 μM with a best-fit Hill coefficient, *n*, of 1.2 (Fig. 2F).

In addition to the speed of vesicle fusion, we analyzed the delay from the onset of the UV-illumination to the onset of the rise of membrane capacitance, which was a free parameter in our fitting functions (see eq. 1). The delay was strongly dependent on the post-flash Ca^2+^ concentration and the dose-response curve showed no signs of saturation at high Ca^2+^ concentrations (Fig. 2G), which is consistent with the non-saturating release rates. These data reveal that the fusion kinetics of synaptic vesicles increased up to a Ca^2+^ concentration of 50 μM without signs of saturation, suggesting a surprisingly low apparent affinity of the fusion sensor at mature cMFBs under physiological temperature conditions (*K_D_* > 50 μM).

**Figure 2.**
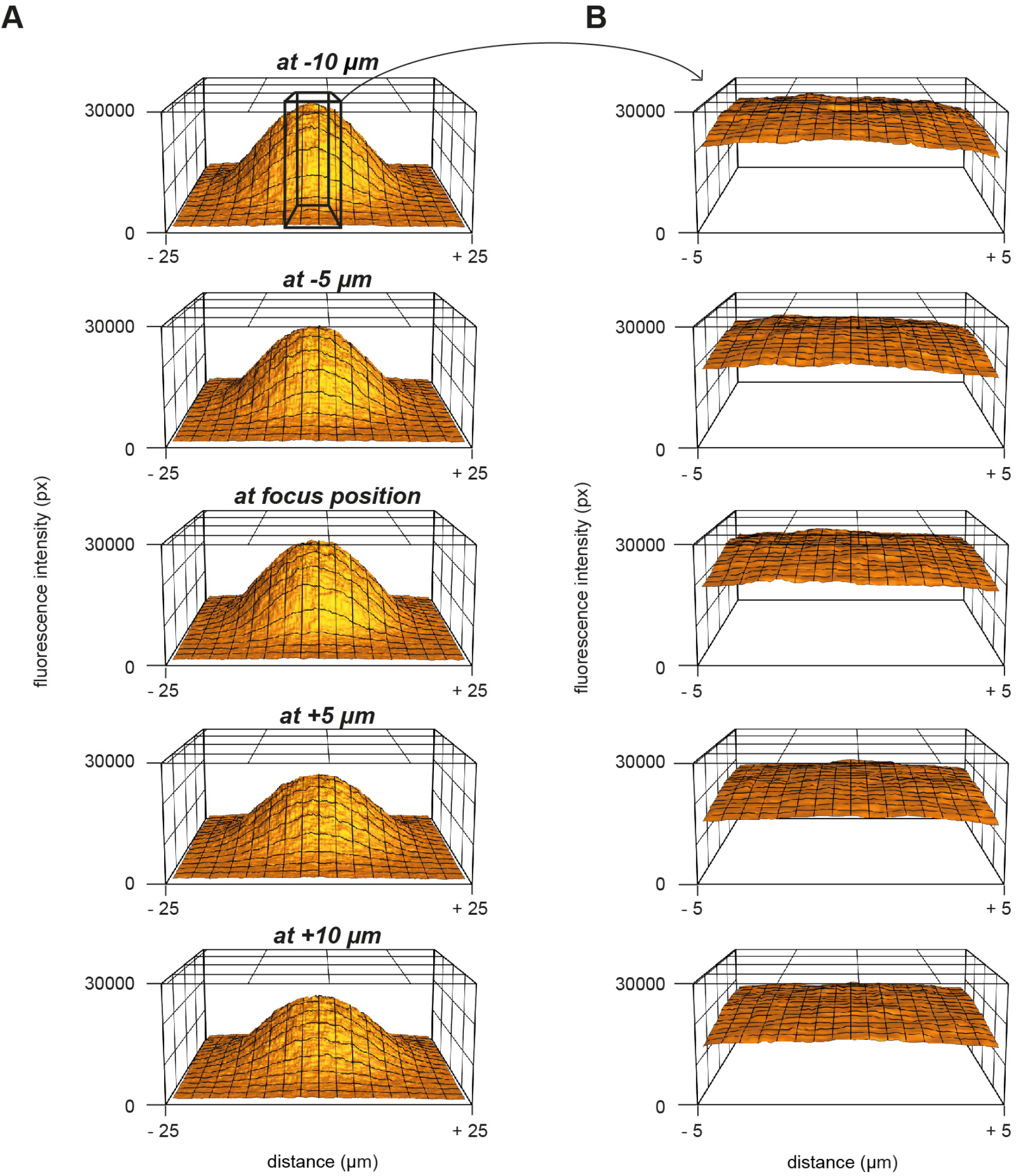
figure supplement 1 Measurement of the UV energy profile with caged fluorescein. A. 3D plot of the fluorescence profile in response to UV uncaging of caged-fluorescein at different z-positions. B. Magnification of the middle part in panel (A) over a range of 10 μm.

### Ca^2+^ uncaging dose-response curve measured with deconvolution of EPSCs

Capacitance recordings are not very sensitive in detecting low release rates. We therefore performed simultaneous pre- and postsynaptic recordings and used established deconvolution techniques to calculate the presynaptic release rate by analyzing the EPSC as previously applied at this synapse (Fig. 3A, B; Ritzau-Jost et al., 2014). Kynurenic acid (2 mM) and cyclothiazide (100 μM) were added to the extracellular solution in order to prevent the saturation and desensitization of postsynaptic AMPA receptors, respectively. Ca^2+^ uncaging in the presynaptic terminal evoked EPSCs with kinetics which strongly depended on the post-flash Ca^2+^ concentration. The cumulative release obtained from deconvolution analysis of the recorded EPSCs was fitted as the capacitance traces (eq. 1). At low Ca^2+^ concentrations (<5 μM), a significant amount of neurotransmitter release could be measured, which is consistent with previous reports from central synapses (Bollmann et al., 2000; Fukaya et al., 2021; Sakaba, 2008; Schneggenburger and Neher, 2000). The presynaptic release rates increased with increasing post-flash Ca^2+^ concentration and no saturation in the release rate occurred in the dose-response curve (Fig. 3D). The dose-response curve for the delay from the onset of the UV illumination to the onset of the rise of the cumulative release trace (eq. 1) did not show signs of saturation of the release kinetics in the investigated range. Thus, consistent with capacitance measurements, deconvolution analysis of postsynaptic currents revealed a shallow Ca^2+^-dependence of neurotransmitter release kinetics (Fig. 3D and E). Fitting a Hill equation to the deconvolution data resulted in a best-fit *K_D_* >50 μM and a Hill coefficient of 1.6 (Fig. 3D). Therefore, two independent measures of synaptic release (presynaptic capacitance measurements and postsynaptic deconvolution analysis) indicate a non-saturating shallow dose-response curve up to ~50 μM.

To rule out methodical errors that might influence the dose-response curve, we carefully determined the *K_D_* of the Ca^2+^ indicator OGB-5N using several independent approaches including direct potentiometry (Fig. 3 – figure supplement 1), because this value influences the estimate of the Ca^2+^ affinity of the fusion sensors linearly. We estimated a *K_D_* of OGB-5N of ~30 μM being at the lower range of previous estimates ranging from 20 to 180 μM (Delvendahl et al., 2015; Digregorio and Vergara, 1997; Neef et al., 2018), arguing against an erroneously high *K_D_* of the Ca^2+^ indicator as a cause for the non-saturation.

In addition, we used the two following independent approaches to rule out a previously described Ca^2+^ overshoot immediately following the UV illumination. Such Ca^2+^ overshoot would be too fast to be detected by the Ca^2+^ indicators (Bollmann et al., 2000) but could trigger strong release with weak UV illumination which, would predict a shallow dose-response curve. First, the time course of Ca^2+^ release from DMn was simulated (see below; Fig. 6A) and no significant overshoots were observed (see below; Fig. 6A). Secondly, we experimentally compared strong and short UV illumination (100% intensity; 0.1 ms) with weak and long UV illumination (10% intensity; 1 ms), because a Ca^2+^ overshoot is expected to primarily occur with strong and short UV illumination. Comparison of these two groups of UV illumination resulted in similar post-flash concentrations but did not reveal a significant difference in the corresponding release rate indicating that undetectable Ca^2+^ overshoots did not affect the measured release rate (Fig. 3 – figure supplement 3). Therefore, both approaches argue against a Ca^2+^ overshoot as an explanation for the shallow dose-response curve.

**Figure 3.**
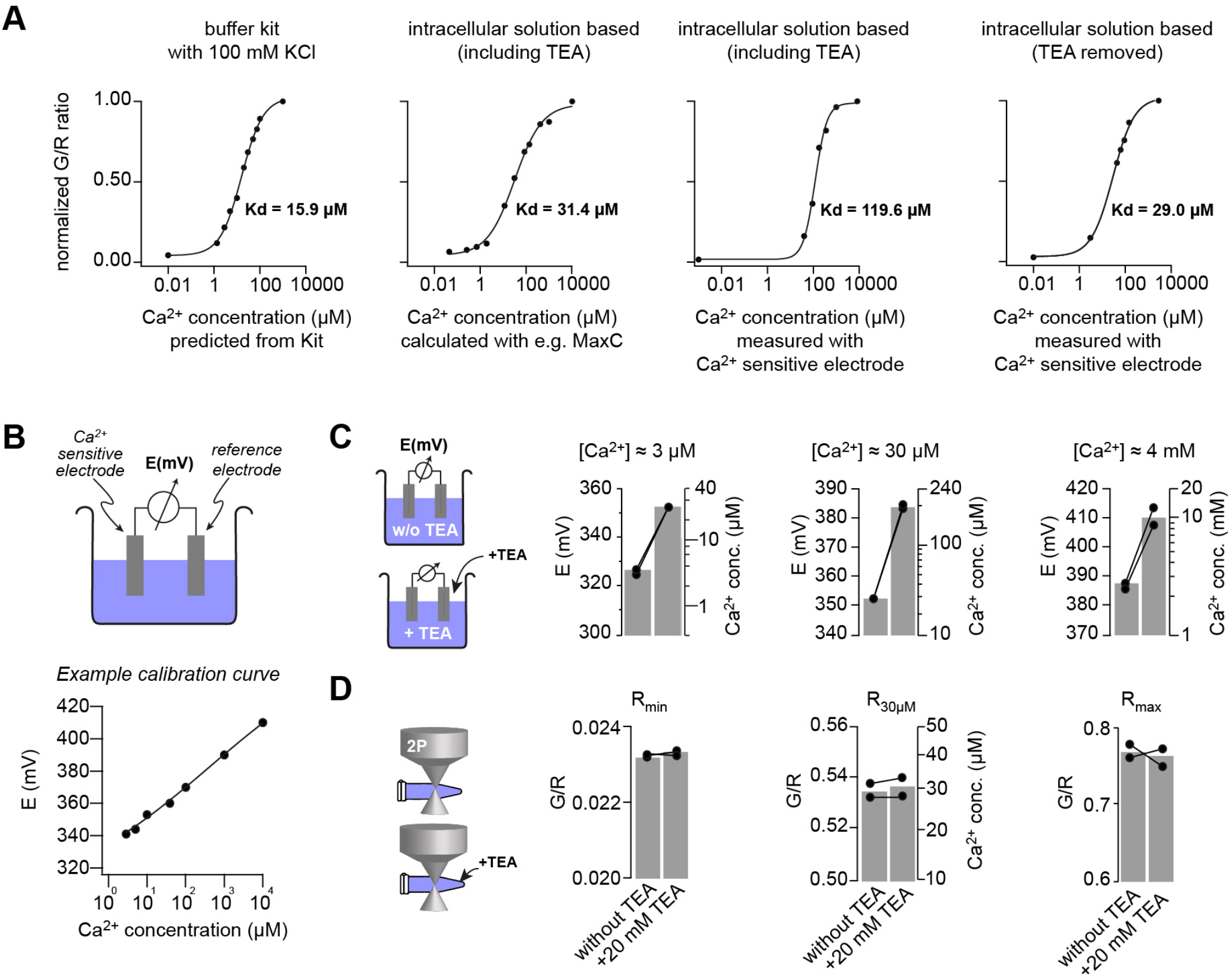
figure supplement 1 Measuring the *K_D_* of the Ca^2+^ sensitive dyes. A. Green (OGB-5N) over red (Atto594) fluorescence ratio for different Ca^2+^ concentrations, measured using either a Ca^2+^ calibration buffered kit or by clamping the free Ca^2+^ using EGTA in the intracellular patching solution. The free Ca^2+^ concentration was predicted from the kit, calculated with software like Maxchelator (MaxC) or measured by potentiometry using a Ca^2+^-sensitive electrode. The indicated *K_D_* values were obtained from superimposed fits with Hill equations. B. *Top:* illustration of the Ca^2+^-sensitive electrode. *Bottom:* Example of a calibration curve of the Ca^2+^-bsensitive electrode fitted with a straight line. C. Effect of Tetraethylammonium (TEA) on the Ca^2+^ sensitive electrode at different Ca^2+^ concentrations. 20 mM TEA induced ~10-fold increase in the potential (left axis) and thus the read-out Ca^2+^ concentration (right axis) of intracellular solutions which had free Ca^2+^ concentrations clamped by EGTA to 3 μM, 30 μM, or 4 mM (pH was kept constant; bargraphs represent the mean; line-connected circles represent two independent repetitions). D. Effect of TEA on G/R fluorescence ratio. The ratio of the intracellular solution containing only 10 mM EGTA (R_min_), free Ca^2+^ clamped with EGTA to 30 μM (R30μM), or 10 mM Ca^2+^ (R_max_) did not change upon adding 20 mM TEA indicating that TEA is not contaminated with Ca^2+^ but instead TEA specifically interferes with the Ca^2+^-sensitive electrode.

**Figure 3.**
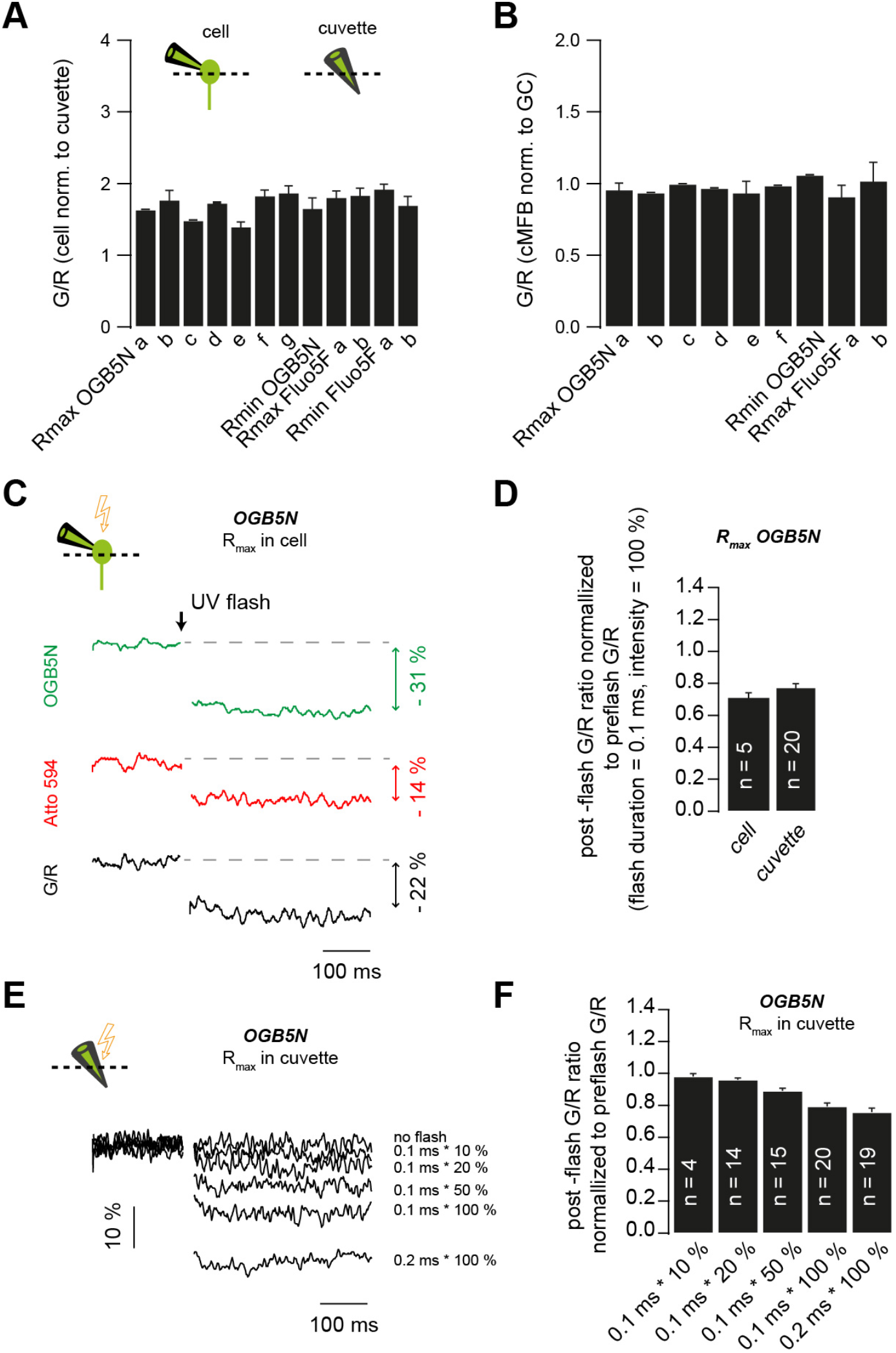
figure supplement 2 Correction for the post-flash changes in the fluorescent properties of the intracellular solution. A. Green over red fluorescence (G/R) ratios measured *in situ* normalized to G/R ratios measured in cuvettes. Data represent the different solutions used throughout the study. (a-g) represent measurements obtained from different solutions prepared using different pre-stocks of the fluorescent indicators or a different DMn/Ca^2+^ concentration. B. Green over red fluorescence (G/R) ratios measured in cMFBs normalized to G/R ratios measured in GCs. Data represent different solutions used throughout the study. (a-f) represent measurements obtained from different solutions prepared using different pre-stocks of the fluorescent indicators or a different DMn/Ca^2+^ concentration. C. Example traces of in situ post-flash alterations in the green fluorescence, in the red fluorescence, and the overall drop in the G/R ratio (in black) in response to a UV flash of 0.1 ms duration and 100 % intensity. D. Comparison of the UV-flash-induced bleaching of fluorescent indicators measured in cells to the UV-flash-induced bleaching of fluorescent indicators measured in cuvettes, in response to a UV flash of 0.1 ms duration and 100 % intensity. E. Example traces of UV-flash-induced changes occurring in cuvettes in response to UV flashes of different intensities or duration. F. Average UV-flash-induced changes occurring in cuvettes in response to UV flashes of different intensities or duration.

**Figure 3.**
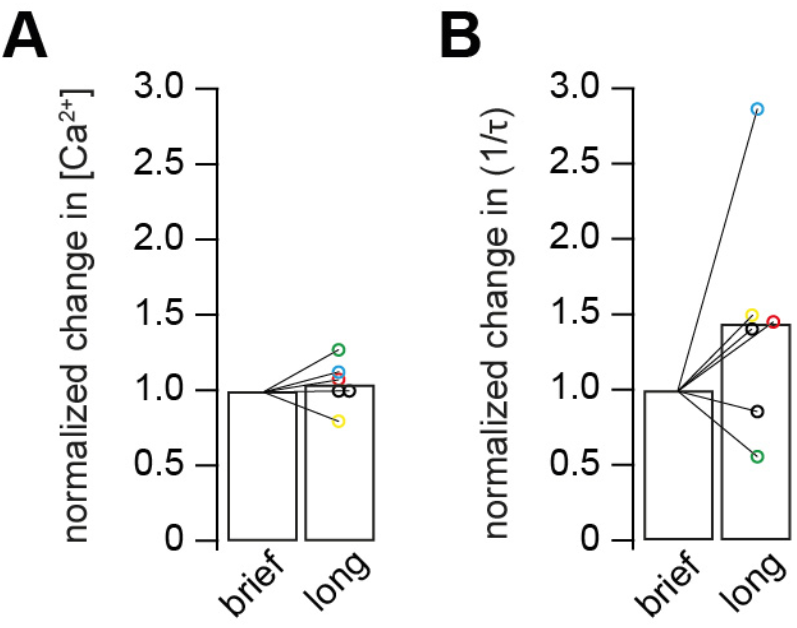
figure supplement 3 Comparison of brief versus long UV illumination to rule out fast Ca^2+^ overshoots. A. Post-flash Ca^2+^ concentration obtained from long flashes of 1 ms duration and 10% UV intensity, normalized to post-flash Ca^2+^ concentration obtained from brief flashes of 0.1 ms duration and 100% UV intensity. B. Release rates obtained from long flashes of 1 ms duration and 10% UV intensity, normalized to release rates obtained from brief flashes of 0.1 ms duration and 100% UV intensity. Color code matches the data in A and B.

### Presynaptic and postsynaptic measurements reveal two kinetic processes of neurotransmitter release

In some Ca^2+^ uncaging experiments, synaptic release appeared to have two components, which could be due to heterogeneity amongst release-ready vesicles. We therefore systematically compared mono- and bi-exponential fits to the capacitance and deconvolution data (Fig. 4 A and B). Several criteria were used to justify a bi-exponential fit (see methods). One criterion was at least a 4% increase in the quality of bi-compared with mono-exponential fits as measured by the sum of squared differences between the fit and the experimental data (χ^2^; Fig. 4D). Consistent with a visual impression, this standardized procedure resulted in the classification of ~40% of all recordings as biexponential (38 out of 80 capacitance measurements and 17 out of 59 deconvolution experiments; Fig. 4C and D). The release rate of the fast component (1/τι) of the merged capacitance and deconvolution data showed no signs of saturation consistent with our previous analyses of each data set separately. Fitting a Hill equation to the merged data indicated a *K_D_* >50 μM and a Hill coefficient of 1.6 (Fig. 4C). The release rate of the slow component (1/τ_2_; if existing) was on average more than 10 times smaller (black symbols, Fig. 4C). These data indicate that there are at least two distinct kinetic steps contributing to release within the first 10 ms.

### Fast and Ca^2+^-independent sustained release

To gain more insights into the mechanisms of sustained vesicle release, we focused on the synaptic release within the first 100 ms after Ca^2+^ uncaging. To investigate the Ca^2+^-dependence of sustained release, we estimated the number of vesicles (Nv) released between 10 and 100 ms after flash onset, assuming a single vesicle capacitance of 70 aF and 90 granule cells-contacts per mossy fiber rosette (see methods; Ritzau-Jost et al., 2014). There was considerable variability in the release rate between 10 and 100 ms, which could be due to differences in bouton size and wash-out of proteins during whole-cell recordings. However, the release rate showed no obvious dependence on the post-flash Ca^2+^ concentration (Fig. 5B). These data indicate that the slope of the sustained component of release is Ca^2+^-independent in the investigated Ca^2+^ concentration range of 1-50 μM, consistent with previously observed Ca^2+^-independent vesicle recruitment as assessed by depolarizing cMFBs to 0mV in the presence of EGTA (Ritzau-Jost et al., 2014).

### Release schemes with five Ca^2+^ steps and fast recruitment via parallel or sequential models can explain Ca^2+^-dependence of release

To investigate mechanisms that could explain a non-saturating and shallow dose-response curve and rapid sustained release, we performed modeling with various release schemes. First, we simulated the exact time course of the concentration of free Ca^2+^. The Ca^2+^ release from DMn and subsequent binding to other buffers and the Ca^2+^ indicator were simulated based on previously described binding and unbinding rates (Faas et al., 2005; Faas et al., 2007; Fig. 6A; see methods). In contrast to previous results, which predicted a significant overshoot of Ca^2+^ following UV illumination with short laser pulses (Bollmann et al., 2000), our simulations predict little overshoot compared to the Ca^2+^ concentration measured by the Ca^2+^ indicator (Fig. 6B). The discrepancy is readily described by recent improvements in the quantification of Ca^2+^ binding and unbinding kinetics (Faas et al., 2005; Faas et al., 2007). The calculations predict an almost step-like increase in the free Ca^2+^ concentration with a 10-90% rise time below 50 μs. These simulated UV illumination-induced transients of free Ca^2+^ concentrations were subsequently used to drive the release schemes. Realistic noise was added to the resulting simulated cumulative release rate and the analysis using exponential fits (eq. 1) was performed as with the experimental data (Fig. 6C).

We compared three different release schemes in their ability to reproduce our experimental data. In model 1, a single pool of vesicles with two Ca^2+^ binding steps was used as previously established, e.g., for chromaffin cells and rod photoreceptors (Duncan et al., 2010; Voets, 2000). Such an assumption would readily explain the shallow dose-response curve (Bornschein and Schmidt, 2018). The two components of release could be replicated by assuming rapid vesicle recruitment from a reserve pool (V_R_; Fig. 6D). However, adjusting the free parameters did not allow reproducing the synaptic delay (Fig. 6E). We therefore tested two more sophisticated models in which vesicle fusion is triggered via five Ca^2+^ binding steps (Schneggenburger and Neher, 2000). In model 2, the first vesicle pool represents the docked vesicles and the second pool represents a replacement pool, which can undergo rapid docking and fusion (Miki et al., 2016; Miki et al., 2018), therefore representing two kinetic steps occurring in sequence. In model 3, two pools of vesicles with different Ca^2+^-sensitivity exist, where both types of vesicles can fuse with different Ca^2+^ affinity (Voets, 2000; Walter et al., 2013; Wölfel et al., 2007), therefore representing two kinetic steps occurring in parallel. Model 3 reproduced the data as good as model 2, however the non-saturation up to 50 μM could be reproduced somewhat better in model 3. Interestingly, models 2 and 3 both replicated the observed shallow dose-response curve despite the presence of five Ca^2+^ binding steps. These results indicate that established models with five Ca^2+^-steps incorporating fast vesicle recruitment via sequential or parallel vesicle pools can replicate our data fairly well.

### Ca^2+^ uncaging with different pre-flash Ca^2+^ concentrations indicates Camdependent vesicle priming

Finally, we aimed to obtain a mechanistic understanding that could explain both the strong dependence of action potential-evoked release on basal Ca^2+^ concentration (cf. Fig. 1) and the Ca^2+^-dependence of vesicle fusion (cf. Figs. 2–6). In principle, the action potential-evoked data in Fig. 1 could be explained by an acceleration of vesicle fusion kinetics or, alternatively, an increase in the number of release-ready vesicles upon elevated basal Ca^2+^. To differentiate between these two mechanistic possibilities, we investigated the effect of basal Ca^2+^ concentration preceding the UV illumination (pre-flash Ca^2+^) on flash-evoked release. The pre-flash Ca^2+^ concentration can only be reliably determined with the Ca^2+^ indicator Flou5F used in the experiments with weak flashes (see Supplementary Table 1). We therefore grouped the deconvolution experiments with weak flashes, which elevated the Ca^2+^ concentration to less than 5 μM, into two equally sized groups of low and high pre-flash Ca^2+^ (below and above a value of 200 nM, respectively). Due to the presence of the Ca^2+^ loaded DMn cage, the pre-flash Ca^2+^ concentrations were on average higher than the resting Ca^2+^ concentration in physiological conditions of around 50 nM (Delvendahl et al., 2015). In both groups, the post-flash Ca^2+^ concentration was on average similar (~3 μM; Fig. 7B). The peak EPSC amplitude of postsynaptic current was significantly larger with high compared to low pre-flash Ca^2+^ concentration (38 ± 10 and 91 ± 16 pA, n = 18 and 13, respectively, P_Mann-Whitney_ = 0.001; Fig. 7A and C). Correspondingly, the amplitude of the fast component of release as measured from deconvolution analysis was larger with high compared to low pre-flash Ca^2+^ (18 ± 5 and 49 ± 10, n = 18 and 13, respectively, P_Mann-Whitney_ = 0.005; Fig. 7C). However, the kinetics of vesicle fusion, measured as the inverse of the time constant of the fast component of release, were not significantly different for both conditions (0.15 ± 0.04 and 0.12 ± 0.03 ms^-1^ for the low and high pre-flash Ca^2+^ conditions, n = 18 and 13, respectively, P_Mann-Whitney_ = 0.74; Fig. 7C). The delay was also not significantly different (P_Mann-Whitney_ = 0.54; Fig. 7C). These data indicate that the number of release-ready vesicles were increased upon elevated basal Ca^2+^ concentration but the fusion kinetics were unaltered. We therefore added an additional Ca^2+^-dependent maturation step to the initial vesicle priming of the release schemes (see methods; note that this was already present in the above-described simulations of Fig. 6 but it has little impact on these data). This allowed replicating the threefold increase in the action potential-evoked release when driving the release scheme with a previously estimated local Ca^2+^ concentration during an action potential (Fig. 7D; Delvendahl et al., 2015). Thus, the release schemes can explain the Ca^2+^-dependence of the recruitment, priming, and fusion of vesicles at mature cMFBs at physiological temperature.

## Discussion

Here, we provided insights into the Ca^2+^-dependence of vesicle recruitment, priming, and fusion at cMFBs. The results obtained at this synapse show prominent Ca^2+^-dependent priming steps, a shallow non-saturating dose-response curve up to 50 μM, and Ca^2+^-mindependent sustained vesicle recruitment. Our computational analysis indicates that the peculiar dose-response curve can be explained by well-established release schemes having five Ca^2+^ steps and rapid vesicle recruitment via sequential or parallel vesicle pools. Thus, we established quantitative scheme of synaptic release for a mature high-fidelity synapse, exhibiting both high- and low-affinity Ca^2+^ sensors.

### Ca^2+^ affinity of the vesicle fusion sensor

The Ca^2+^-sensitivity of vesicle fusion seems to be synapse-specific. In contrast to the estimated Ca^2+^ affinity for vesicle fusion of ~100 μM at the bipolar cell of goldfish (Heidelberger et al., 1994) and the squid giant synapse (Adler et al., 1991; Llinás et al., 1992), recent studies showed that the affinity is much higher at three types of mammalian central synapses: the calyx of Held (Bollmann et al., 2000; Lou et al., 2005; Schneggenburger and Neher, 2000; Sun et al., 2007; Wang et al., 2008), the inhibitory cerebellar basket cell to Purkinje cell synapse (Sakaba, 2008), and the hippocampal mossy fiber boutons (Fukaya et al., 2021). Consistent with reports from mammalian central synapses, our data revealed prominent vesicle fusion at concentrations below 5 μM arguing for a high-affinity fusion sensor (Figs. 2–4). However, the non-saturation of the dose-response curve (Figs. 2–4) argues for the presence of a rather low-affinity fusion sensor at cMFBs. In our simulations, both model 2 and 3 exhibit vesicles with a Ca^2+^-affinity similar to the calyx of Held. Nevertheless, with high intracellular Ca^2+^ concentrations (>20 μM) these vesicles will fuse very rapidly and the further increase in the release kinetics (causing the non-saturating dose-response curve) can be explained by rapid vesicle recruitment from a sequential pool of vesicles exhibiting use-dependent lowering of the Ca^2+^-affinity (V1 in model 2; Miki et al., 2018) or from a parallel pool of vesicles with lower Ca^2+^ affinity (V_1_ in model 3; Hallermann et al., 2010). Our data therefore indicate that the shallow and non-saturating dose-response curve is the consequence of rapid recruitment of vesicles that still exhibit a lower Ca^2+^-affinity compared to fully recovered vesicles. Consistent with this interpretation, a lowering in the Ca^2+^-affinity of the vesicle fusion sensor has been observed at the calyx of Held with Ca^2+^ uncaging following vesicle depletion (Müller et al., 2010; Wadel et al., 2007). These newly recruited vesicles might contribute particularly to the dose-response curve at the cMFB because the cMFB has a much faster rate of vesicle recruitment compared with the calyx of Held synapse (Miki et al., 2020) providing a possible explanation why the here-reported dose-response curve differs from previous results at the calyx of Held. Furthermore, cMFBs seem to have functional similarities with ribbon-type synapses because it has recently been shown that the vesicle mobility in cMFBs is comparable to ribbon-type synapses (Rothman et al., 2016). The hallmark of ribbon-type synapses is their rapid vesicle recruitment (Lenzi and von Gersdorff, 2001; Matthews, 2000) and indeed more shallow dose-response curves were obtained at the ribbon photoreceptors and inner hair cell synapses (Duncan et al., 2010; Heil and Neubauer, 2010; Johnson et al., 2010; Thoreson et al., 2004), but see (Beutner et al., 2001). Therefore, these results predict similar shallow non-saturating dose-response at other central synapses with rapid vesicle recruitment (Doussau et al., 2017; Miki et al., 2016; Pulido and Marty, 2017).

### Ca^2+^-sensitivity of vesicle priming

The steps preceding the fusion of synaptic vesicles are in general still poorly understood (Südhof, 2013). There is evidence that some steps preceding the fusion are strongly Ca^2+^-mdependent (Neher and Sakaba, 2008), as has been demonstrated at chromaffin cells (Voets, 2000; Walter et al., 2013) and at several types of synapses such as the calyx of Held (Awatramani et al., 2005; Hosoi et al., 2007), the crayfish neuromuscular junctions (Pan and Zucker, 2009), parallel fiber to molecular layer interneuron synapses (Malagon et al., 2020), and cultured hippocampal neurons (Chang et al., 2018; Stevens and Wesseling, 1998). In previous reports, the Ca^2+^-dependence of vesicle priming at cMFBs was analyzed more indirectly with the Ca^2+^ chelator EGTA (Ritzau-Jost et al., 2014; Ritzau-Jost et al., 2018) and the obtained results could be explained by Ca^2+^-dependent models but surprisingly also by Ca^2+^-independent models (Hallermann et al., 2010; Ritzau-Jost et al., 2018). Furthermore, the analysis of molecular pathways showed that the recovery from depression is independent of the Ca^2+^/calmodulin/Munc13 pathway at cMFBs (Ritzau-Jost et al., 2018). Our paired recordings and uncaging experiments (Figs. 1 and 7) clearly demonstrate pronounced Ca^2+^-dependence of vesicle priming at cMFBs. Taken together, these data indicate that some priming steps are mediated by Ca^2+^mdependent mechanisms, which do not involve the Ca^2+^/calmodulin/Munc13 pathway. A potential candidate for such a Ca^2+^-dependent mechanism are the interaction of diacylgylcerol/phospholipase C or Ca^2+^/phospholipids with Munc13s (Lee et al., 2013; Lou et al., 2008; Rhee et al., 2002; Shin et al., 2010).

Here, we used single action potentials (Fig. 1) and weak uncaging stimuli (post-flash Ca^2+^ concentration of ~3 μM; Fig. 7) to investigate the impact of the basal Ca^2+^ concentration. Synaptic vesicles that fuse upon single action potentials and weak uncaging stimuli are particularly fusogenic and thus might represent the superprimed vesicles with a particular high release probability (Hanse and Gustafsson, 2001; Ishiyama et al., 2014; Kusch et al., 2018; Lee et al., 2013; Schlüter et al., 2006; Taschenberger et al., 2016) suggesting that the process of superpriming is Ca^2+^-dependent. This interpretation would also provide an explanation why in a recent report, triggering an action potential in the range of 10-50 ms time before another action potential (which elevates basal Ca^2+^ concentrations) restored the synchronicity of synaptic vesicle fusion in mutant synapses which has a phenotype of synchronous-release-impairment (Chang et al., 2018). It would be furthermore consistent with a proposed rapid, dynamic, and Ca^2+^-dependent equilibrium between primed and superprimed vesicles (Neher and Brose, 2018). However, further investigations are needed for the dissection between the Ca^2+^mdependence of priming and superpriming. Yet, our data show that some priming steps are strongly Ca^2+^-dependent with a high-affinity Ca^2+^ sensor that allow detecting changes between 30 and 180 nM at cMFBs.

### Ca^2+^-sensitivity of vesicle recruitment

The upstream steps of vesicle priming, referred to as recruitment, refilling, or reloading, remain controversial in particular with respect to their speed. The slow component of release (during prolonged depolarizations or Ca^2+^ elevations with uncaging) was initially interpreted as a sub-pool of release-ready vesicles that fuse with slower kinetics (see e.g. Sakaba and Neher, 2001). However, recent studies indicate very fast vesicle recruitment steps (Blanchard et al., 2020; Chang et al., 2018; Doussau et al., 2017; Hallermann et al., 2010; Lee et al., 2012; Malagon et al., 2020; Miki et al., 2016; Miki et al., 2018; Saviane and Silver, 2006; Valera et al., 2012). These findings further complicate the dissection between fusion, priming, and recruitment steps. Therefore, the differentiation between ‘parallel’ release schemes with fast and slowly fusing vesicles and ‘sequential’ release schemes with fast vesicle recruitment and subsequent fusion is technically challenging at central synapses. Our data could be described by both sequential and parallel release schemes (model 2 and 3; Fig. 6). The non-saturation of the release rate could be described somewhat better by the parallel model 3. However, further adjustment of the use-dependent slowing of the rates in model 2 (see *k*_on,plug_, *k*_off,plug_, and eq. 3 and 4; Miki et al., 2018) can result in a sequential model exhibiting both fast and slowly fusing vesicles with different Ca^2+^-sensitivity (see Mahfooz et al., 2016, for an alternative description of use-dependence of vesicle fusion). Such use-dependent sequential models ultimately complicate the semantic definitions of ‘sequential’ and ‘parallel’, because the newly recruited vesicles will fuse in a molecularly different state, which could also be viewed as a parallel pathway to reach fusion. Independent of the difficulty to differentiate between sequential and parallel release schemes, the sustained component of release exhibited little calcium dependence in the here-tested range between 1 and 50 μM (Fig. 5). The Ca^2+^-independence of vesicle recruitment in the investigated range is consistent with the previously observed EGTA-independent slope of the sustained release during prolonged depolarizations (Ritzau-Jost et al., 2014). Our data cannot differentiate if recruitment is mediated by a fully saturated Ca^2+^ sensor for priming (mode 2; assumed Kd of 2 μM; Miki et al., 2018) or a parallel Ca^2+^-independent step (mode 3). Thus, during sustained activity at cMFBs vesicle recruitment is either mediated by fully Ca^2+^-independent processes or by an apparently Ca^2+^-independent processes in the relevant Ca^2+^ concentration range because of a saturated high-affinity Ca^2+^ sensor.

### Mechanistic and functional implications

The Ca^2+^-sensitivity of vesicle fusion critically impacts the estimates of the coupling distance between Ca^2+^ channels and synaptic vesicles, mainly those obtained based on functional approaches (Neher, 1998; Eggermann et al., 2011; but not on structural approaches, see e.g. Éltes et al., 2017; Rebola et al., 2019). Our previous estimate of the coupling distance at the cMFB of 20 nm (Delvendahl et al., 2015) was based on the release scheme of Wang et al. (2008) obtained at the calyx of Held synapse at an age of (P16-P19) at room temperature and assuming a Q10 factor of 2.5. The now estimated *kon* and *K_off_* rates at mature cMFBs at physiological temperature were slightly larger and smaller than the temperature-corrected values from the calyx, respectively, resulting in a slightly higher affinity of the fast releasing vesicles (V2 in model 2 and 3). Therefore, at the cMFB, the coupling distance of the vesicles released by a single action potential is if anything even smaller than the previous estimate of 20 nm.

In addition, our data might provide a link between Ca^2+^-dependent priming and facilitation. Synaptotagmin-7 is a high-affinity Ca^2+^ sensor (Sugita et al., 2002) that could mediate the here-reported three-fold increase in synaptic strength (Figs. 1 and 7). Synaptotagmin-7 has been proposed to play a role in synaptic facilitation at different synapses supporting a molecularly distinct mechanism of facilitation (Jackman and Regehr, 2017). An increase in the size of the fusogenic sub-pool of release-ready vesicles mediated by basal Ca^2+^ might provide the underlying mechanism where Synaptotagmin-7 could be a sensor for the changes in basal Ca^2+^ levels and therefore affect synaptic strength (Liu et al., 2014).

Finally, synaptic fidelity has been shown to increase with age at cMFBs (Cathala et al., 2003), neocortical synapses (Bornschein et al., 2019), and the calyx of Held (Fedchyshyn and Wang, 2005; Nakamura et al., 2015; Taschenberger and von Gersdorff, 2000).

During high-frequency transmission, the residual Ca^2+^ concentration increases up to a few μM at cMFBs (Delvendahl et al., 2015) but mature cMFBs can still sustain synchronous release (Hallermann et al., 2010; Saviane and Silver, 2006). The developmental decrease in the affinity of the release sensors observed at the calyx of Held (Wang et al., 2008) and the here-reported shallow-dose-response curve at mature cMFBs could be an evolutionary adaption of synapses to prevent the depletion of the release-ready vesicles at medium Ca^2+^ concentrations and therefore allow maintaining sustained synchronous neurotransmission with high fidelity (Matthews, 2000).

## Acknowledgement

We thank Erwin Neher for help with algorithms for calculating the Ca^2+^ concentration of the intracellular solutions (Fig. 1) and for helpful discussions. This work was supported by a European Research Council Consolidator Grant (ERC CoG 865634) to S.H and by the German Research Foundation (DFG; SCHM1838/2) to H.S.

